# Identification of cognate recombination directionality factors for large serine recombinases by virtual pulldown

**DOI:** 10.1101/2024.06.11.598349

**Authors:** Heewhan Shin, Alexandria Holland, Abdulrazak Alsaleh, Alyssa D. Retiz, Ying Z. Pigli, Oluwateniola T. Taiwo-Aiyerin, Tania Peña Reyes, Adebayo J. Bello, Femi J. Olorunniji, Phoebe A. Rice

## Abstract

Integrases from the “large serine” family are simple, highly directional site-specific DNA recombinases that have great promise as synthetic biology and genome editing tools. Integrative recombination (mimicking phage or mobile element insertion) requires only integrase and two short (∼40 – 50) DNA sites. The reverse reaction, excisive recombination, does not occur until it is triggered by the presence of a second protein termed a Recombination Directionality Factor (RDF), which binds specifically to its cognate integrase. Identification of RDFs has been hampered due to their lack of sequence conservation and lack of synteny with the phage integrase gene. Here we use Alphafold2-multimer to identify putative RDFs for more than half of a test set of 98 large serine recombinases, and experimental methods to verify predicted RDFs for 4 of 5 integrases chosen as test cases. We find no universally conserved structural motifs among known and predicted RDFs, yet they are all predicted to bind a similar location on their cognate integrase, suggesting convergent evolution of function. Our methodology greatly expands the available genetic toolkit of cognate integrase – RDF pairs.

## INTRODUCTION

Large serine recombinases (LSRs), which are encoded by many lysogenic bacteriophages (and some other mobile genetic elements), have great potential as genetic tools.(1–5)(6–9)(10)(11) In their natural setting, this family of site-specific DNA recombinases catalyzes unidirectional site-specific recombination between an ∼50bp bacteriophage (“phage”) attachment site (*attP*) and an ∼40bp bacterial site (*attB*), resulting in the insertion of the phage genome into the host chromosome (Figure 1a). The resulting prophage can then be passively replicated as part of the bacterial chromosome.(12) Upon activation of the phage’s lytic phase, a second phage-encoded protein, the Recombination Directionality Factor (RDF), binds the integrase protein and alters its preferred reaction direction to greatly favor the excision reaction. Although LSRs are readily identified in genomic sequences due to their conserved sequence motifs, RDFs are not, and the relatively few that are known were identified primarily through painstaking genetic work. (13–15)(16–18)(19–21) (22)(23)(24)(25). No conserved sequence motifs are apparent among them, nor do they show consistent synteny with their cognate integrase genes. Here we describe and experimentally verify a new Alphafold2-multimer-based method for rapid identification of RDFs – essentially, a virtual pulldown approach. (26–28)

**Figure 1.**
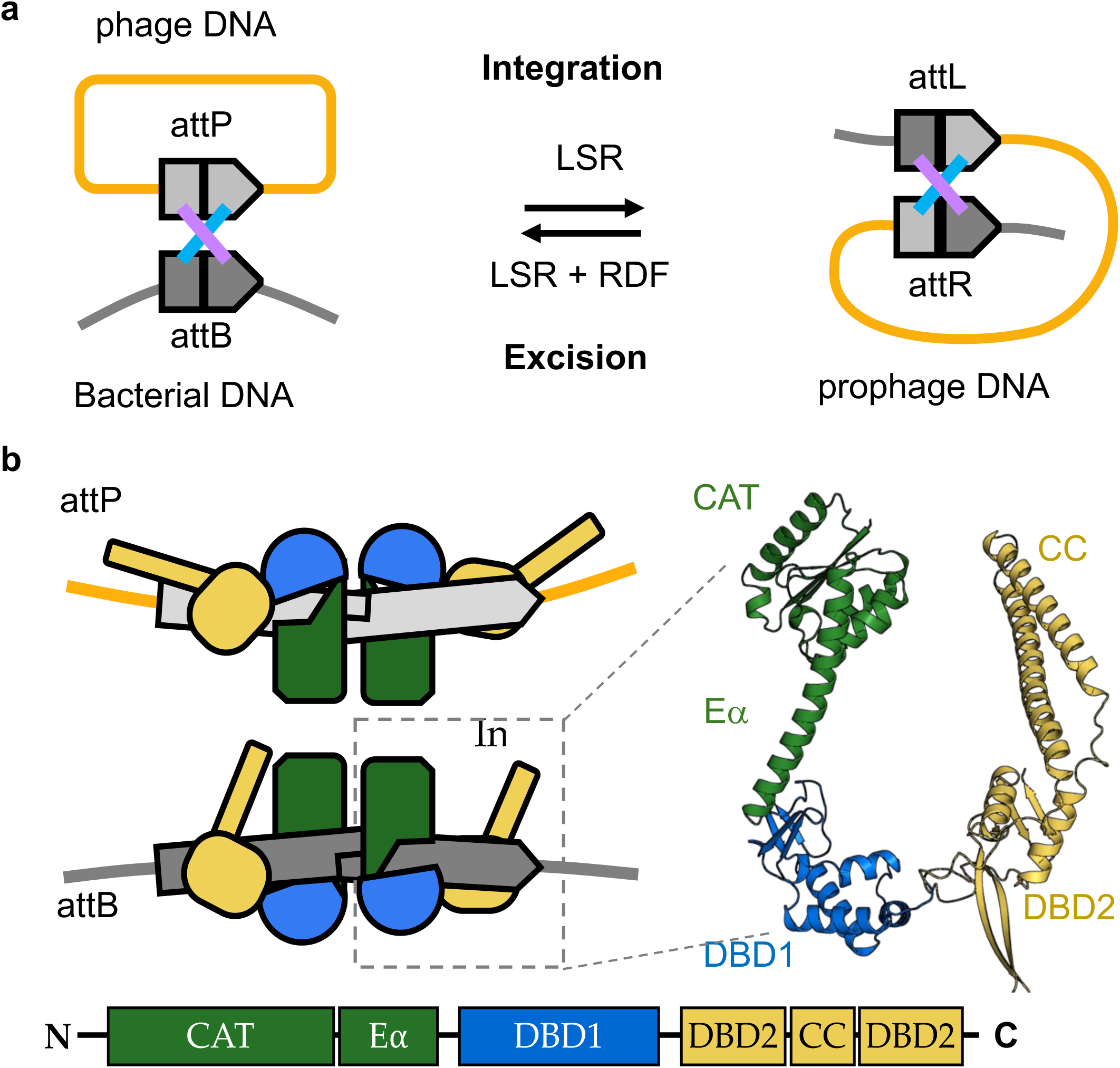
Recombination Mechanism and Domain Structure. a) Integrative recombination requires only the LSR and its cognate *att* sites. The reverse reaction (excisive recombination) requires the presence of a second protein, the cognate RDF. b) Domain structure of LSIs. Left panel: cartoon of LSR dimers bount to *attP* (top) and *attB* (bottom) sites. Right panel: predicted structure of A118 integrase. Bottom: linear cartoon of LSI domain organization. CAT: catalytic domain; Ea: long helix connecting the CAT to the DNA-binding domains; DBD1: DNA-binding domain 1 (also known as recombinase domain); DBD2: DNA-binding domain 2 (also known as Zn-binding domain); CC: coiled coil.

Many features of LSRs render them useful as genetic tools. Unlike CRISPR-Cas systems, they have evolved to catalyze the insertion of large payloads, and their recombination mechanism leaves the DNA product with not a single broken phosphodiester bond. In contrast to bacteriophage integrases from the mechanistically very different tyrosine family, they do not require host proteins and their *attP* sites are much smaller.(29) Although LSRs lack the programmable sequence specificity of RNA-guided systems, their lack of target flexibility can be alleviated by a “drag-and-drop” procedure that uses a CRISPR-Cas – derived system to insert an LSR’s *attB* site, or by choosing from the large array of already characterized LSRs with differing sequence specificity.(9, 10, 30, 31) This toolkit was greatly expanded by the recent publication of a list of over 60 LSRs that can catalyze insertions into the human genome. (10) Furthermore, for some applications, such as the SIRA method for assembly of large replicons, the availability of multiple LSRs with orthogonal sequence specificity is a key feature that facilitates multiplexing. (32)(33) When the cognate RDFs are known for particular LSRs, their versatility as tools expands significantly. For example, RDF-mediated reaction reversal can be used for modular editing of assembled replicons, and LSR-RDF pairs can be used to create living logic gates.(6)

LSRs constitute a branch of the “serine” family of site-specific DNA recombinases, which share a conserved overall mechanism of recombination (Figure 1a).(34) Each crossover site in the DNA is bound by a dimer of the recombinase, after which the two DNA-bound dimers are brought together by a regulatory apparatus that varies widely among systems but is an intrinsic part of the crossover site-bound protein subunits in the LSR case. The serine nucleophile in the active site of each subunit then attacks a particular phosphodiester bond, displacing a 3’ OH to create a reaction intermediate in which both DNA duplexes have double-strand breaks with 2-nt 3’ overhangs, and each 5’ end is covalently linked to a recombinase subunit. The recombinase tetramer then swivels internally to realign the broken ends, and the DNAs are religated by the reverse of the DNA cleavage reaction. The reaction direction is determined not by a change in chemical bond energy but by energetic differences among the substrate and product conformations of the protein-DNA complexes.

Serine recombinases have modular domain organizations. The “large” family, which includes LSRs, carries the conserved catalytic domain at its N-terminus, followed by two DNA binding domains (DBDs). For simplicity, we refer to these as DBD1 and DBD2, although DBD1 is often termed “recombinase domain” and DBD2 “zinc-binding domain.” The integration reaction is thought to be driven forward by a change in the protein-protein interactions mediated by the hydrophobic tip of a coiled-coil that is inserted within DBD2. (Figure 1b; see reference (35) for details of the proposed mechanism). Biochemical studies have found that RDFs bind to the coiled-coil and/or DBD2 of their cognate integrases, presumably reversing the preferred reaction direction by altering the range of trajectories available to the coiled coil.(19)(22)(23)

A detailed understanding of how LSRs and their cognate RDFs function has been hampered by a lack of structural information. Only very partial structures are available for LSRs: the isolated catalytic domains, the two DBDs bound to half of an *attP* site, and isolated coiled-coil dimers.(35–37) However, recent AI-based advances in protein structure prediction can now help to fill these knowledge gaps.(26, 27) We found that AlphaFold2-multimer predicts a similar binding site at the DBD2-coiled coil junction for known LSR-RDF pairs, despite the lack of any conserved structural motif among those RDFs.

We then automated Alphafold2-multimer to efficiently predict RDFs for LSRs whose RDFs were previously unknown, using the large collections of active LSRs identified by Durrant et al. (10) and Yang et al. (31) as test cases. Finally, we show that RDFs for 4 of the 5 integrases picked for *in vivo* verification do indeed function as predicted. We further verified the function of two of these pairs *in vitro* using purified proteins. The virtual pulldown workflow described here will lead to identification of the RDFs for many known integrases and for those yet to be discovered. This will give access to a larger pool of integrase-RDF pairs available for fundamental studies on the reaction mechanism of the integrase-RDF system, and for building orthogonal integrase-based genetic circuits for synthetic biology applications.

## MATERIAL AND METHODS

### Virtual pulldowns

#### Preparation of paired input files for AlphaFold2 multimer predictions

We used the collections of LSRs organized by Durrant *et al*. and Yang *et al.* as test cases for our procedure. (10)(31) For each LSR listed in their supplementary tables, we obtained the corresponding genomic data (DNA and protein coding sequences) from the National Center for Biotechnology Information (NCBI) database. We utilized the first 15-20 and last 15-20 nucleotides of the *attB* and *attP* sequences to search within the genome sequence and identify the prophage region, and in some cases corroborated that range using Phaster.(38)

The protein coding sequences within the identified prophage region were extracted as prey sequences. For the bait sequence, we predicted the LSR’s structure with ESMFold (https://esmatlas.com/resources?action=fold; (39)) and then truncated the sequence to include only the second DNA-binding domain (DBD2). The truncated “bait” LSR sequence was paired with each prey sequence in a FASTA format for AlphaFold2-multimer predictions. In some cases where the initial pulldown failed to produce a clear “hit”, it was repeated using both DNA binding domains or the intact LSR. Python scripts used for this and later steps can be found as Supplementary Data.

#### Multimer predictions

LocalColabFold v.1.5.2 was installed on Beagle-3, a shared GPU cluster at the University of Chicago’s Research Computing Center (RCC), following the steps described in https://github.com/YoshitakaMo/localcolabfold. (26, 27, 40) We note that the power of the virtual pulldown approach was independently reported by Yu et al. (28) while we were optimizing our method and experimentally testing our results.

#### Assessment of output files

LocalColabFold generates Predicted Aligned Error (PAE) plots with predicted scores and models (in JSON and PDB format, respectively). For efficient assessment, all PAE plots were concatenated; pTM and ipTM scores were extracted from JSON files, and these values were plotted using gnuplot. Predicted models exhibiting low PAE and high pTM and ipTM scores were visually evaluated in PyMOL. Initial hits were chosen as those prey proteins that gave the highest ipTM value, which usually exceeded the pTM value. Potential hits were rejected if they (1) were not predicted to interact with DBD2, (2) were predicted to interact with the tip of the coiled coil, as that hydrophobic patch is known to be required for self interactions, or (3) were predicted to interact with the DNA-binding surface of DBD2. (36) The latter proteins were most likely DNA mimics involved in the host-virus arms race, and suggests that virtual pulldowns may provide a new way to discover such proteins. (41)

#### Structure figures

All structure figures were made using PyMOL (The PyMOL Molecular Graphics System, Version 2.0 Schrödinger, LLC.; https://pymol.org/).

### Testing of Predictions

To experimentally test our predictions, we chose 5 examples: Nm60, Bt24, and Cb16 from the Durrant *et al*. list as well as Int10 and Int 30 from the Yang *<u>et al</u>*. list. (10, 31) Candidate RDFs were tested using a slight variation of the *in vivo* inversion assay that we and others have previously used to study integrase recombination reactions *in vivo* (42–44). (See diagram in Figure 6). Each assay requires two co-transformed plasmids: (1) a test plasmid carrying the *att* sites for the integrase in question (in inverted orientation) flanking a promoter that drives expression of either GFP or RFP, depending on its orientation, and (2) an expression vector for the integrase or integrase – RDF fusion of interest. The activity of two integrase – RDF pairs (Nm60 and Int30) and were additionally verified in *in vitro* assays.

#### Vectors encoding integrase and integrase-RDF fusion proteins

Coding sequences for all integrases and integrase -RDF fusions were cloned between NdeI and XhoI sites in pBAD33, which carries an arabinose-inducible promoter, a p15a origin, and chloramphenicol resistance. The use of an inducible expression system prevented potential issues with toxicity of constitutively expressed integrases, and the use of integrase – RDF fusions (previously described for known integrase – RDF pairs (45)) alleviated potential issues with the stoichiometry of the two proteins that could occur if they were expressed separately. In the fusion constructs, each integrase and its putative RDF were covalently joined together using an 18-residue linker (TSGSGGSGGSGGSGRSGT) between the C-terminal residue of the integrase and the second amino acid residue in the candidate RDF. The amino acid sequences of the proteins of interest were reverse-translated and the DNA sequences codon-optimised for expression in *E. coli* and ordered as gene fragments from Twist Biosciences. Each integrase gene has a SpeI site before the stop codon, which adds the 1^st^ two amino acids (TS) of the linker sequence to the C-terminus of the protein. RDF genes with the linker added to their N-termini were then cloned in-frame between the SpeI and XhoI sites in the integrase vectors.

All new plasmids were verified by sequencing.

#### Prediction of core attP and attB recombination sites

To determine the *attP* and *attB* sequences for integrases of interest, we used the information provided by Durrant *et al*. and Yang *et al.* (10, 31) in combination with genomic sequences. We used the half-site sequence alignment approach of Van Duyne & Rutherford to assign potential binding sites for the two DNA binding domains and to determine the most likely location of the central dinucleotide (Figure 5). (46)

#### *In vivo* recombination reactions

*In vivo* recombination assays (inversion) of each integrase - putative RDF pair were carried out using the invertible promoter reporter system depicted in Figure 6A. The plasmid substrates were based on the previously reported pϕC31-invPB and pϕC31-invRL ((42); see Figures 4 and 6). The backbone of these vectors carries a pSC101 origin and a kanamycin resistance marker that are compatible with those on our protein expression vectors. The recombination sites (*attP* and *attB*) for each integrase were cloned into pϕC31-invPB to replace the *att* sites for ϕC31 integrase using gene fragments (Twist Biosciences) covering the entire ∼560bp invertible segment. Substrate plasmids containing *attL* and *attR* for each integrase were made by integrase – mediated recombination.

*E. coli* DS941 strains (47) containing the substrate plasmid (kanamycin selection) and the arabinose-inducible expressing vector (chloramphenicol) for each integrase or integrase-RDF fusion were prepared in advance such that recombination activities can be initiated when integrase expression is induced by addition of arabinose to a growing culture. To prepare the recombination strain, competent *E. coli* DS941 cells were transformed with the inversion substrate plasmid and the integrase expression vector and grown for 16 hours in LB media in the presence of kanamycin (50 µg/ml) and chloramphenicol (25 µg/ml) selection.

To assay *attP* x *attB* recombination activity of each integrase, fresh overnight culture of each recombination strain was grown at 37 C in LB media to mid-log phase in the presence of kanamycin (50 µg/ml) and chloramphenicol (25 µg/ml). Integrase or integrase-RDF expression was induced for 2 hours by addition of 0.2% arabinose after which cultures were diluted 1:1000 in fresh LB media containing 0.2% glucose to repress further integrase expression and grown for 16 hours with shaking at 37 C. Cultures were diluted 1/100, 000 in fresh LB media and spread on LB agar plates containing kanamycin (50 µg/ml). Slight variations were used in assaying Int10: cells were grown in 0.2% glucose before switching to arabinose for induction, induction was continued overnight, and DNA from the overnight culture was miniprepped and retransformed before plating.

Recombination of *attP* and *attB* sites inverts the orientation of the promoter, thereby switching gene expression from RFP to GFP (Figure 6B), and generating *attR* and *attL* sites. Colonies expressing GFP or RFP were imaged in a Gel Doc™ XR+ imaging system (Bio-Rad, Hercules, CA, USA) using blue epi illumination and a 530 nm / 28 nm bandpass filter for GFP or green epi illumination with a 605 nm / 50 nm bandpass filter for RFP. Exposure times were 0.01s and 0.02s respectively. Variations for Int10 were: red epi illumination with a 695 nm / 55 nm bandpass filter for RFP, and exposure times of 0.025s and 0.25s, respectively.

Following recombination, individual colonies expressing GFP were grown in liquid culture for 16 hours with kanamycin selection. Plasmid DNAs were extracted from the liquid culture and separated on 1.0% agarose gel. Bands corresponding to supercoiled substrate (inversion products bearing *attR* x *attL* sites) were cut out of the gel, purified, and used to retransform *E. coli* cells. Plasmid DNA samples were subsequently extracted from the cultures and sequenced to verify that they contain the expected *attR* and *attL* sites for each integrase. The verified recombination products of *attP* x *attB* reactions (*attR* x *attL* plasmid substrates) were then used for the experiments where the activities of the candidate RDFs were tested. In those assays, recombination of *attR* x *attL* sites leads to inversion of the promoter resulting in expression of RFP, while stopping GFP production (Figure 6).

#### Protein expression and purification

The expression and purification of Nm60 integrase and Nm60 integrase-RDF fusion was as described previously (45). The DNA sequences encoding the integrase and the integrase-RDF fusion were cloned between NdeI and XhoI sites in pET28a(+) and the plasmids were used to transform *E. coli* BL21(DE3)pLysS strain. The strain for each protein was grown at 37 ^◦^C in LB to an optical density 0.8, before cooling the cultures to 20 ^◦^C and protein expression induced with 0.5 mM IPTG. Protein expression was allowed to continue for 16 hours at 20 ^◦^C. Each protein was purified by nickel affinity chromatography and bound proteins were eluted with an imidazole gradient buffer system. Samples of fractions corresponding to peaks were analysed on SDS-PAGE and chosen fractions containing the desired proteins were dialysed against Protein Dilution Buffer, PDB (25 mM Tris–HCl (pH 7.5), 1 mM DTT, 1 M NaCl and 50% glycerol), and stored at –20 ^◦^C. Dilutions of the integrase and the integrase-RDF fusion were made into the same buffer for *in vitro* recombination reactions.

#### *in vitro* recombination reactions

Recombination assays of excisive (*attR* x *attL*) and integrative (*attP* x *attB*) activities were carried out on the same plasmid substrates used for the *in vivo* reactions (Figure 7A). Plasmid DNA substrates for Nm60 integrase were grown in *E. coli* DS941 and purified using Qiagen miniprep kit. *In vitro* recombination of supercoiled plasmid substrates and analysis of recombination products were carried out using conditions similar to those described previously (33, 43).

Typically, recombination reactions were carried out by adding integrase (0.5-1.0 µM, 5 µl) to a 30 µl solution containing the plasmid substrate (25 µg/ml), 50 mM Tris–HCl (pH 7.5), 100 µg/ml BSA, 5 mM spermidine, and 0.1 mM EDTA. Reactions were allowed to proceed at 30 ^◦^C for 2 hours, after which the integrases were denatured by heating the reaction at 80 ^◦^C for 10 minutes to stop all recombination activities. The samples were cooled and treated with the restriction endonuclease XhoI (New England Biolabs) to facilitate analysis of recombination products. Following the restriction endonuclease treatment, the reaction products were treated with 0.1% SDS and protease K before reaction products were separated by 1.2% agarose gel electrophoresis (33, 43).

## RESULTS

### Previously identified RDFs: variable structures bind a conserved LSR feature

We first examined several previously identified LSR – RDF pairs, using Alphafold2-multimer to predict the structures of their complexes. Because protein structure can be more conserved than sequence, we had expected to find a small conserved structural motif within the RDFs that would mediate interaction with their cognate LSRs. Figure 2 shows that we did not: some RDFs are predicted to utilize only alpha helices as interaction motifs (e.g. phiRv1), some to use both helices and beta strands (e.g. Bxb1 and SPbeta), and some to use loops (e.g. phiC31). However, despite sharing no structural homology at all, these disparate RDFs were all predicted to bind DBD2 at or near its junction with the coiled coil. Confidence in these predictions can be seen in the PAE (predicted alignment error) in Supplementary Figure 1. Supplementary Figure 2 shows that these models also agree existing biochemical data. (18, 22, 23)

**Figure 2.**
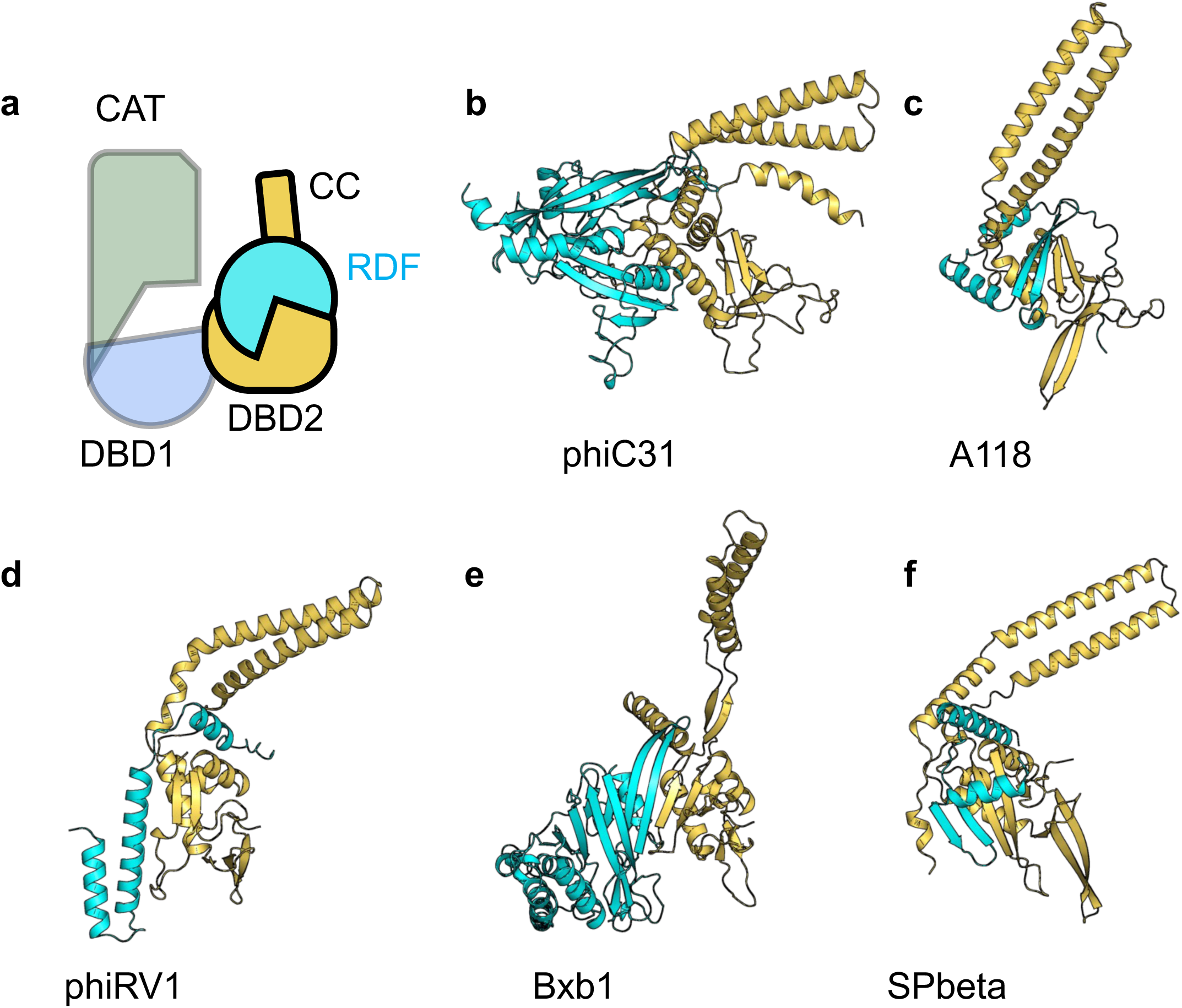
Predicted structures for known LSR-RDF pairs. a) Cartoon similar to those in Figure 1, but with the RDF added (cyan). b-f) Predicted complexes. Only DBD2 and the CC inserted within it (yellow) are shown for the LSR partner but the entire predicted structure structure for each RDF (cyan) is shown.

### Virtual pulldowns identify known RDFs

Given the difficulties in sequence-based RDF prediction and the apparent success of AI-based structure predictions, we asked if Alphafold2-multimer could be used to identify RDFs. The examples shown in Figure 2 were chosen as positive controls. Structures for each ORF within a given phage were predicted in complex with DBD2 of the relevant LSR (see Materials and Methods). For 4 out of these 5 examples, the complex predicted with the highest confidence was indeed that with the previously-identified RDF. Predictions for the 5^th^, phage Bxb1, were ambiguous until the procedure was repeated using both DNA binding domains as “bait” even though the RDF is only predicted to interact with DBD2. For consistency, the virtual pulldowns shown for all 5 examples in Figure 3 and Supplemental Figure 3 were carried out using the last 400 amino acids of the LSR, which ensured including both DNA binding domains in the bait.

**Figure 3.**
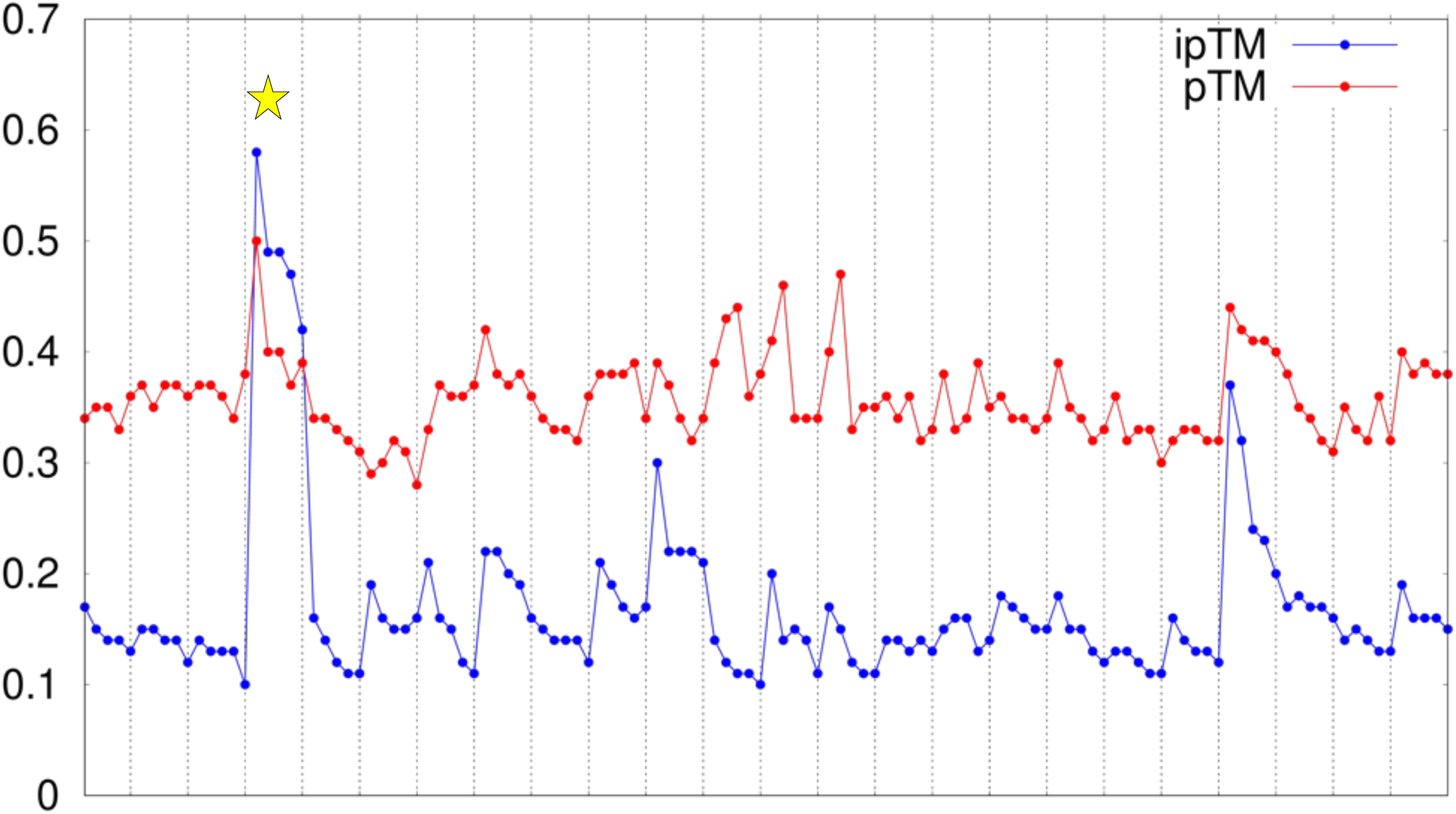
A scatter plot of the Virtual pulldown for the phage-like element A118. Five models were predicted for the complex of each element-encoded protein with the A118 integrase (dotted vertical lines separate sets of models). The plot shows two measures of confidence for each — **p**redicted **T**emplate **M**odeling (pTM; red) and **i**nterface **p**redicted **T**emplate **M**odeling (ipTM; blue) scores — which were calculated using AlphaFold2 multimer v3 model in ColabFold v.1.5.2. The highest-randing model for the known RDF is highlighted with a yellow star. See Supplementary Figure 3 scatter plots for the other LSR-RDF pairs shown in Figure 2 and for the PAE plots for the each of these known LSR-RDF pairs.

We also tested the predicted orthogonality of these LSR – RDF pairs: that is, whether each RDF is predicted to interact with all integrases, or only with its cognate integrase. While some closely-related LSR-RDF pairs do cross-react, e.g. those of the ϕC31 family (23, 43), less closely related RDFs have been shown not to. For example, the RDF for Bxb1 integrase (gp47) does not interact with TG1 and ϕC31 integrases (45). Alphafold2-multimer was indeed able to discern the correct pairings: Among this set, the most confidently complexes were always between the cognate pairs (Supplementary Figure 4).

### Prediction of new RDFs by virtual pulldown

Next we used our virtual pulldown procedure to try to predict RDFs for all of the LSRs listed in Supplementary table 2 of Durrant *et al*. as well as the 34 new LSRs identified by Yang *et al*. (10, 31). Of the 68 listed by Durrant et al., we could not find the appropriate mobile element or prophage range for 3 (Ec06, Kp03, Sa10). In our first round, in which we used only DBD2 of each of the remaining 65 LSRs as prey, we were able to make confident predictions in 33 cases (Supplementary Table 1; supplementary data). In examining the failures, we noted that one integrase (Efs2) was in fact a DDE recombinase (48) not an LSR, and that many have additional structured protein at the C-termini that would be expected to block the RDF binding region noted above. Noting our findings for the Bxb1 test case, we repeated the failed pulldowns using a larger fragment of the integrase protein as prey, resulting in 10 more RDFs predicted with moderate to high confidence.

In total, we were able to predict putative RDFs for 43 of the 64 integrases listed by Durrant et al., and to speculatively predict that at least some of the 24 of the remaining integrases may not in fact have a cognate RDF, in some cases perhaps due to substantial C-terminal extensions of the integrase protein that block the potential RDF-binding site. The predicted complexes chosen for experimental testing are shown in Figure 4, with corresponding PAE plots in Supplementary Figure 5. Many of the LSRs (57 % or 38 out of 67) were from genomic elements that Phaster deemed unlikely to be bacteriophages and may instead be other genomic islands with different control mechanisms. However, some of those LSRs for which we were able to confidently predict an RDF also appeared to be from non-bacteriophage mobile genetic elements. We were also able to predict putative RDFs for many of the 34 integrases by Yang *et al.* (Supplementary Table 1). In some cases, our procedure predicted more than one putative RDF, in which case all are listed.

**Figure 4.**
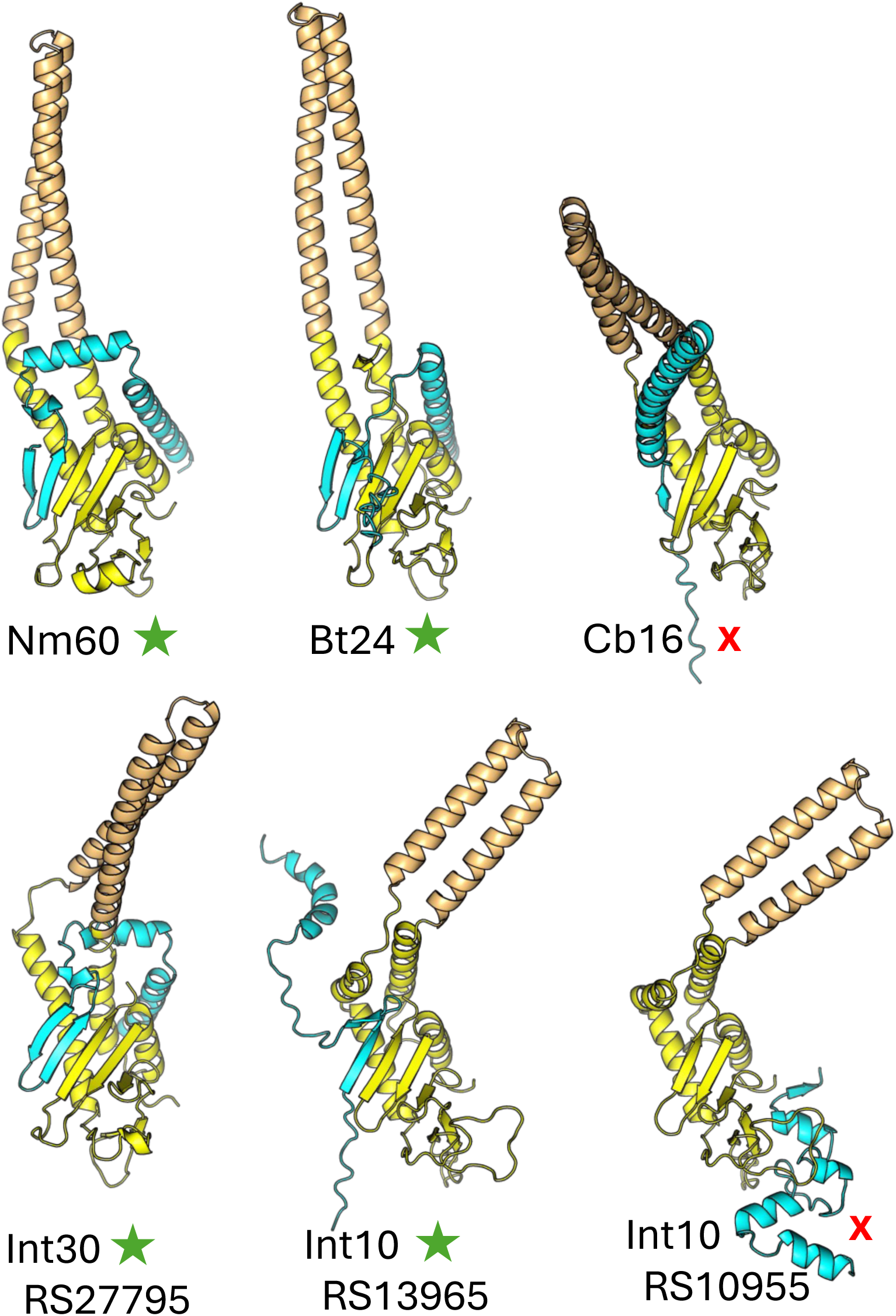
Predicted DBD2-RDF structures complex structures for the 6 LSR – RDF pairs tested experimentally. DBD2 of the LSR is shown in yellow; the CC in tan, and the RDF in cyan. If more than one potential RDF was predicted, a partial locus tag is added to disambiguate. Functional RDFs are marked with green stars and false positives with red x’s. PAE plots for each pair are shown in Supplementary Figure 5.

The predicted RDFs adopt a variety of folds (Figure 4). They are clearly not predicted to fully restrain the mobility of the coiled coil. However, we noted a strong tendency of the RDF to interact not only with DBD2 itself but also with the DBD2-proximal segment of the coiled coil (which is sometimes replaced with beta strands; e.g. see Bxb1 in Figure 2), suggesting that it may partially restrain it.

Finally, we examined the predicted orthogonality for a subset of our collection of newly predicted LSR-RDF pairs (those for which the RDFs were small proteins that were predicted to add beta strands to DBD2’s beta sheet). While some RDFs were exclusively predicted to interact with their cognate LSR, others were predicted to be more promiscuous, although their orthogonality remains to be experimentally tested (Supplementary Figure 6; supplementary data).

### Experimental verification of newly predicted RDFs

To verify our method, we focused on 3 predicted LSR-RDF pairs from the Durrant et al. list (Nm60, Bt24, and Cb16) as well as 2 examples from the Yang et al list (Int10 and Int30). Nm60, Bt24 and Cb16 were the most active integrases shown in Figure 2i of reference (10) for which we could predict a cognate RDF. Int10 was chosen because two potential RDFs were predicted, and we were curious as to whether or not both were active. Two possible RDFs were also predicted for Int30, but only the 1^st^ (RS27795) was tested. All of these predicted RDFs were active except for the Cb16 one and one of the two possibilities for Int10. The function of the RDFs for Nm60 and Int30 were further confirmed *in vitro* using purified proteins. Results for the Nm60 case are shown in the main figures below and results for others are shown in the supplementary figures.

#### Confirmation of Integrase and att site functionality

First, we verified the functionality of our integrase and substrate constructs. Figure 6B (Panel 1) shows the activities of Nm60 integrase and its fusion recombinase on *attP* x *attB* substrate. The results show Nm60 integrase mediated the complete switching of gene expression from RFP to GFP following inverting the orientation of the promoter as a result of *attP* x *attB* recombination reaction (Figure 6A). Similar results were obtained for the other integrases tested (Supplementary Figure 7). This confirms that the 40-50 bp core sequences that we identified are correct and sufficient for *attP* x *attB* recombination.

Following successful *attP* x *attB* recombination by the integrases, we isolated the recombinant DNA plasmid product and used this as *attR* x *attL* substrate for the excisive recombination reaction. The plasmids were sequenced to ascertain *attR* and *attL* sites were formed and to confirm the central dinucleotide crossover point we deduced based on half-site alignment (Figure 5). The sequenced *attR* x *attL* sites for the tested integrases agreed with the predicted crossover points. Finally, we found that, as expected of LSRs, the integrases were essentially inactive on their cognate *attR* x *attL* substrates (Figure 6 and supplementary Figure 7).

**Figure 5:**
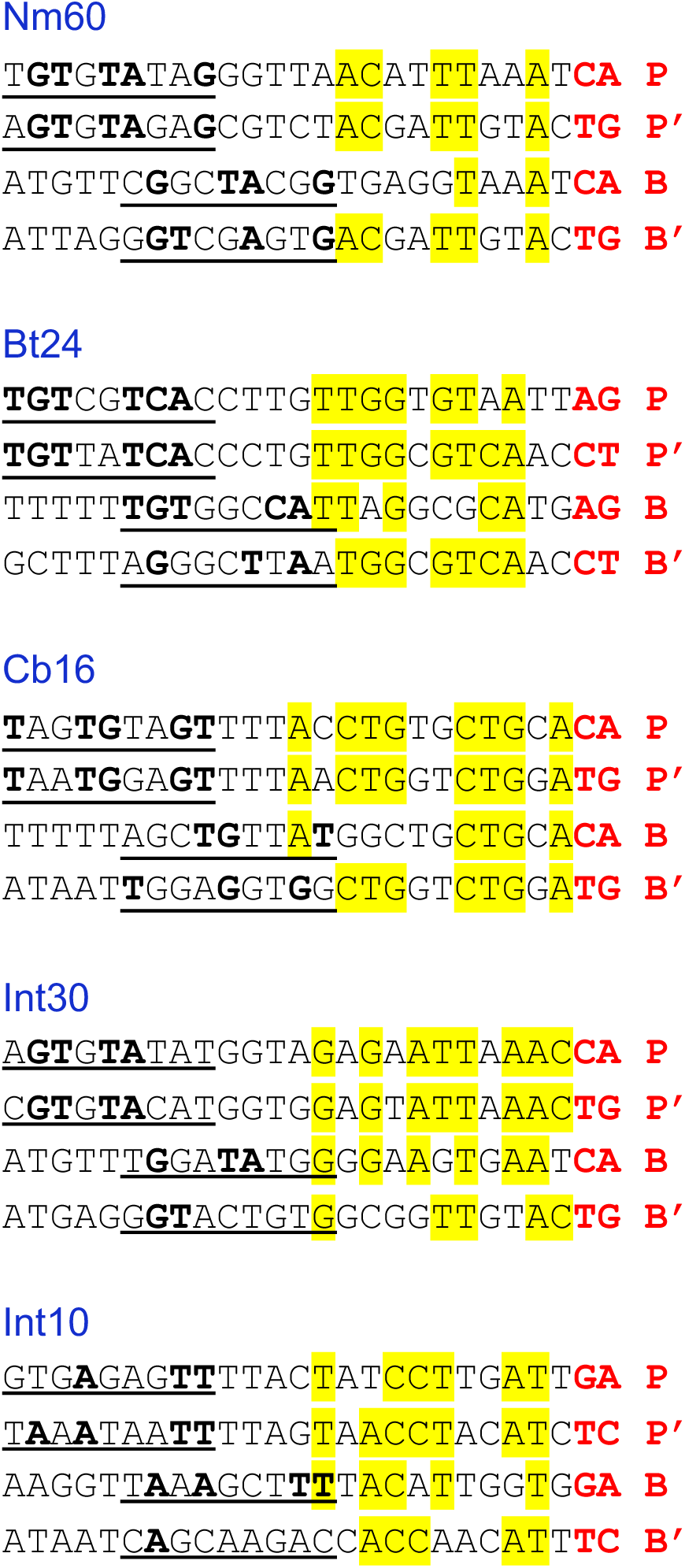
Alignment of LSR half-site sequences. The central dinucleotides are shown in bold and red font. DNA binding domain1 (DBD1, also called “recombinase domain” is expected to bind the innermost 12 bases, and DBD2 (also called “zinc binding domain”) is expected to bind the outer underlined segment. Bases that are conserved in three or four positions are highlighted in yellow in the DBD1 motif and in bold in the DBD2 motif. This convention follows reference (46).

**Figure 6:**
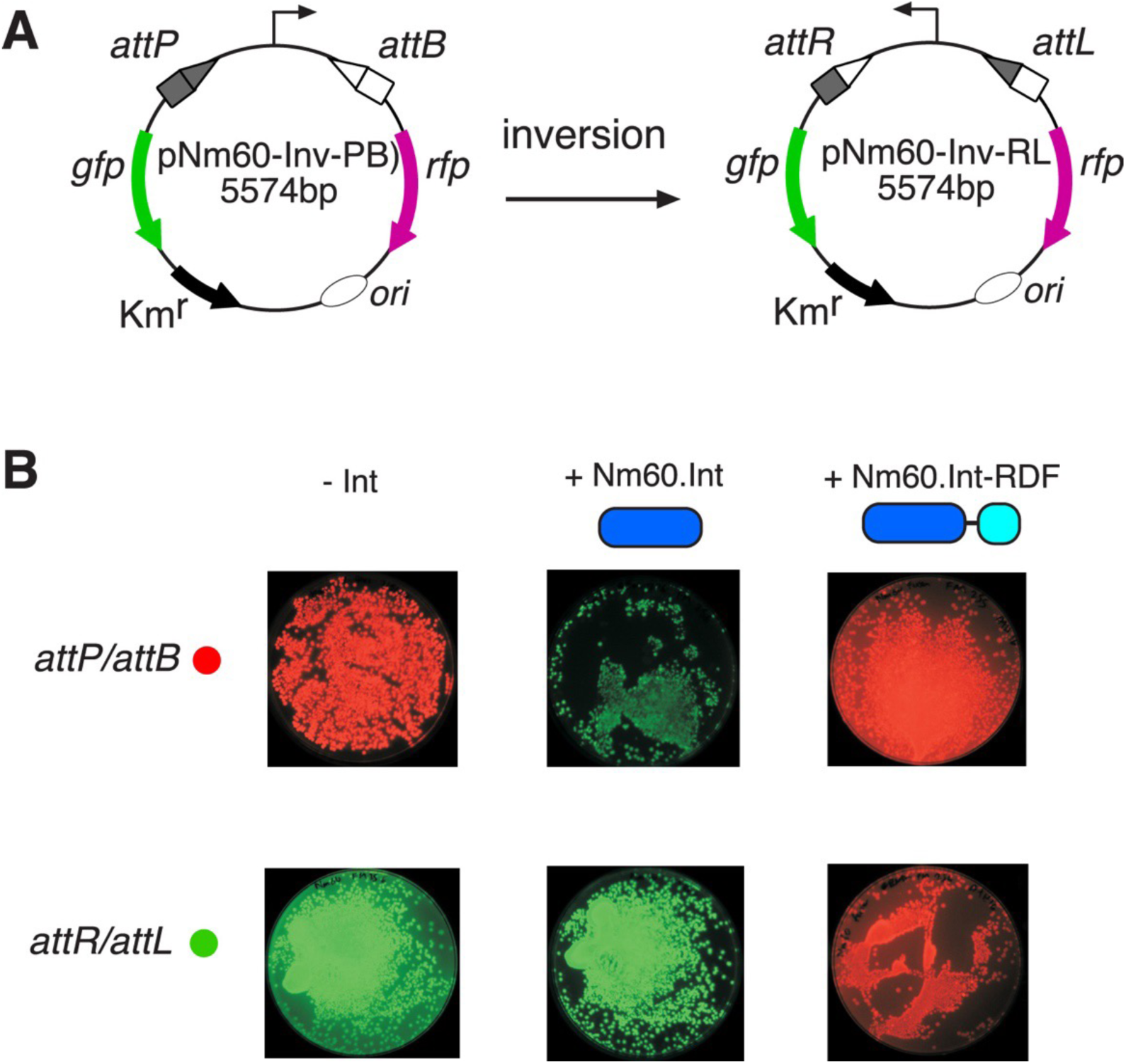
*In vivo* recombination reactions of Nm60 integrase and its fusion with the RDF. a) Schematic illustration of the *in vivo* recombination (inversion) assay. In the plasmid pϕC31-invPB, the promoter, flanked by *attP* and *attB* sites, constitutively drives the expression of a red fluorescent protein (*rfp*) gene (pink arrow). To prevent transcriptional read-through to the green fluorescent protein (*gfp*) gene (green arrow), a terminator sequence is inserted upstream of the promoter. LSR-catalysed recombination (inversion) of *attP* x *attB* to give *attR* x *attL* products (plasmid pϕC31-invRL) flips the orientation of the promoter to allow the expression of GFP, and block RFP production. b) Recombination activities of Nm60 integrase and its fusion to the RDF on *attP* x *attB* (pϕC31-invPB) and *attR* x *attL* (pϕC31-invRL) substrates. The integrase is depicted as a long oval (blue) and the RDF as short a short oval (cyan).

#### In vivo assay of RDF function

A functional RDF should inhibit the *attP* x *attB* recombination by its cognate LSR and activate *attL* x *attR* recombination (12). Figure 6b and Supplementary Figure 7 show that this is indeed the case for the predicted RDFs for Nm60, Bt24, Int10 and Int30. For experiments using these LSR-RDF fusion constructs, the final colonies were predominately red regardless of whether the starting substrate was the *attP* x *attB* one (red) or the *attL* x *attR* one (green). However, the predicted RDF for Cb16 and one of the two predicted Int10 RDFs failed to affect recombination: for these LSR-RDF fusions, the final colonies were predominately green regardless of which starting substrate was used (that is, these fusions behave as if no RDF is present). These results confirm the efficacy of our virtual pulldown approach, with the caveat that two of 6 tested hits tested appear to be false positives.

In all *in vivo* experiments, plasmids were extracted and sequenced to confirm that the *att* sites in the product plasmids are as expected and in agreement with orientation of the promoter directing the expression of GFP or RFP as shown in Figure 6. Results were fully consistent with the predictions in Figure 5.

#### In vitro recombination activities of Nm60 integrase and Nm60 integrase-RDF fusion

Next, we tested the *in vitro* recombination activities of Nm60 integrase and its integrase-RDF fusion in both *attP* x *attB* and *attR* x *attL* reactions. The proteins were expressed and purified as previously described for ϕC31 and Bxb1 integrases and their RDF fusions (45). As shown in Figure 7A, recombination of pϕNm60-invPB (*attP* x *attB*) to give pϕNm60-invRL (*attR* x *attL*) is accompanied by inversion of the DNA segment flanked by the *att* sites. Treatment of the recombination reaction product with the restriction endonuclease, XhoI, gave distinct restriction patterns for recombined and non-recombined DNA.

**Figure 7:**
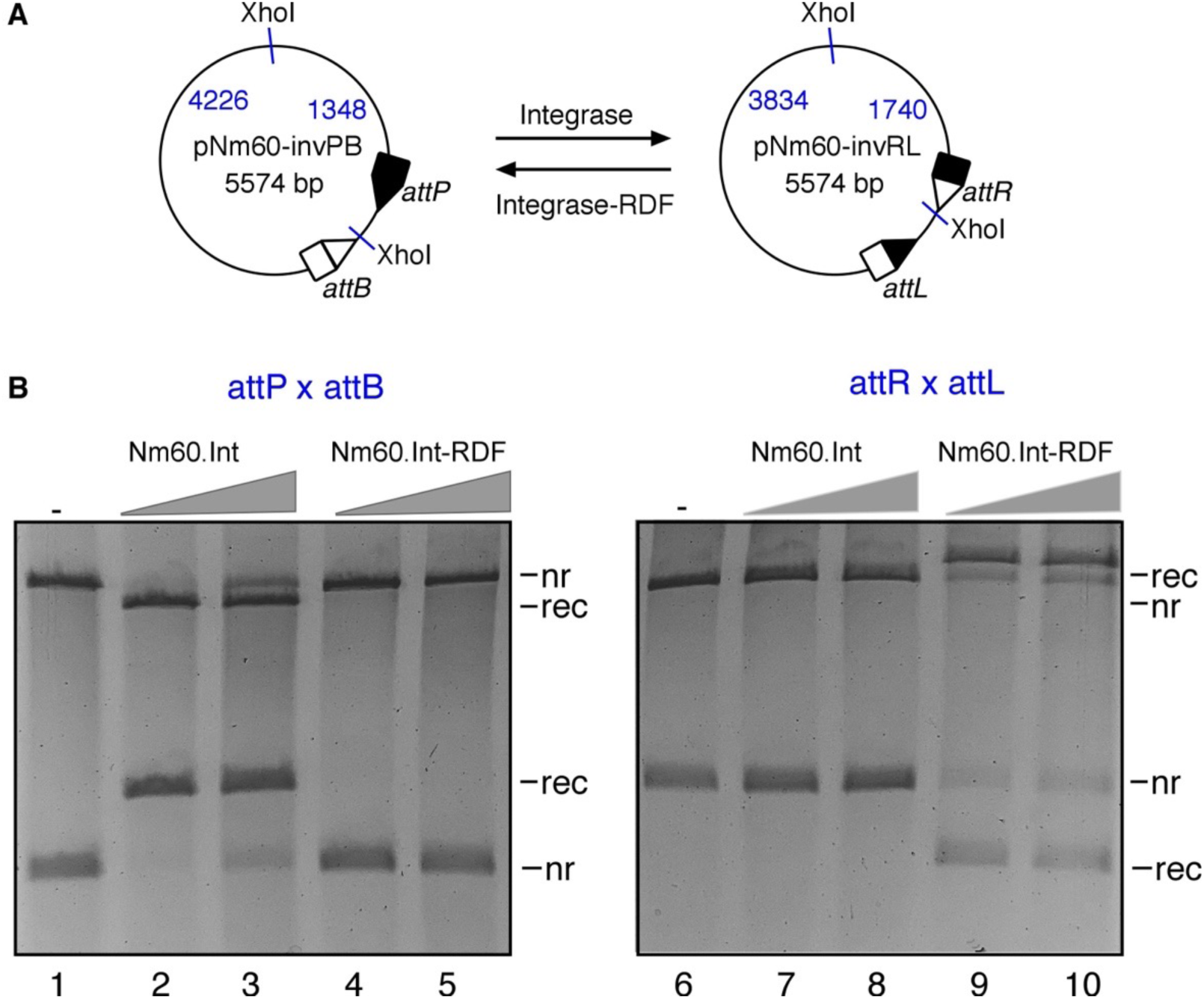
*In vitro* recombination reactions of Nm60 integrase and its fusion with the RDF. a) Schematic illustration of the *in vitro* recombination (inversion) assay. LSR-catalysed recombination (inversion) of *attP* and *attB* sites in pNm60-invPB gives rise to *attR* and *attL* sites in the product plasmid (pNm60-invRL), and vice versa. b) Reactions were carried out for 2 hours as described in Materials and Methods. Integrase and integrase-RDF fusions were used at two final concentrations of 50 and 100 nM. Reaction products were digested with the restriction endonuclease, XhoI, prior to 1.2% agarose gel electrophoresis. As illustrated in (**a**), digestion of the plasmids with XhoI gives different restriction patterns for recombined and non-recombined DNA. The bands on the gel are labeled *nr* (non-recombinant, i.e. substrate), *rec* (recombination product).s

The activities of the two proteins on *attP* x *attB* substrate are shown in Figure 7B (first panel). Nm60 integrase catalysed complete conversion of the substrate, while fusion of the RDF to the integrase completely blocks *attP* x *attB* recombination. The second panel shows the results of the *attR* x *attL* reaction. As expected, purified Nm60 integrase failed to catalyse the reaction, while the integrase-RDF fusion recombined *attR* x *attL* to give *attP* x *attB*. These *in vitro* recombination activities agree with the *in vivo* results described above (Figure 6). Similar results were found for purified Int30 and Int30-RDF fusion proteins (Supplementary Figure 8).

## DISCUSSION

Using Alphafold2-multimer to perform virtual pulldowns, we identified putative RDFs for approximately ∼60 % of the our test set of integrases. (Supplementary Table 1). (10) For initial tests of our predictions, we chose cases for which the integrases were described as highly active by Durrant *et al.* (Nm60, Bt24, and Cb16) as well as two described by Yang *et al.* (Int10 and Int30) (10, 31). For 4 of these 5 integrases, at least one predicted RDF stimulated *attL* x *attR* recombination and inhibited *attP* x *attB* recombination, as expected of an RDF. It is unclear why the Cb16 pulldown returned a false positive. However, in comparing the two potential RDFs tested for Int10, we noted that the non-functional one, while predicted to interact with DBD2, was not predicted to interact with the DBD2-coiled coil junction region as seen for the functional RDFs. This demonstrates the feasibility of our approach, which can obviate the need for painstaking genetic experiments to find the cognate RDF for a given LSR. It also adds to the growing body of reports demonstrating the power of the AI-based virtual pulldown approach (28).

It remains to be determined whether or not the LSRs in our test set for which we did not confidently predict RDFs actually require an RDF to stimulate excisive (*attL* x *attR*) recombination and inhibit integrative (*attP* x *attB*) recombination. Furthermore, although the *attP* x *attB* recombination activity of most of the LSRs in our test set was previously verified (10, 31), their activity on a full range of *att* site pairs remains untested. The LSRs encoded by the staphylococcal SCC*mec* element can catalyze bidirectional recombination on a large variety of *att* site pairings, suggesting that their directionality may be differently regulated, if at all. (49) Alphafold2 models of the SCC*mec* LSRs showed that they do have additional beta strands at the C-terminus of DBD2 that could block the RDF binding site here. However, further studies are needed to determine if the existence of such C-terminal extensions is an accurate predictor of a lack of a cognate RDF and a lack of directionality, or if some LSRs with such extensions have found an alternate solution to directionality. Furthermore, there is precedent for one mobile genetic element to rely on another to supply an excision-stimulating RDF, which we would not have found with our protocol. (25)

### Access to a larger pool of characterised LSR-RDF pairs

The virtual pulldown workflow described here will greatly facilitate identification of the RDFs for known integrases and for those yet to be discovered. This will give access to a larger pool of integrase-RDF pairs available for fundamental studies on the reaction mechanism of the integrase-RDF system. Insights gained from studying several structurally diverse RDFs and how they interact with their cognate integrases could provide a better understanding of the quintessential properties of an RDF. This knowledge could be used to iteratively create a set of structure and property profiles to find common themes among the highly variable RDFs and their LSR interactions, and thus a better understanding of what is important for optimal function.

AlphaFold2-multimer predicts that despite their structural diversity, all the verified RDFs that we looked at are predicted to bind their cognate integrase at the DBD2-coiled coil junction (Figure 2 and 4), while one false positive was predicted to bind elsewhere on DBD2. We envisage that the availability of a wide range of known RDFs will ultimately allow the development of a universal method for designing synthetic RDFs or RDF-independent integrases engineered to catalyse the excisive *attR* x *attL* recombination. Additional structural information will be required to achieve these goals.

### Genetic circuits & logic gates

The potential application of large serine integrases in building genetic circuits and logic gates have been explored in prokaryotic and eukaryotic systems (6, 6, 7, 44, 50–52). These examples have been built using the same set of few characterised LSRs and their RDFs. The new RDFs characterised here and the virtual pulldown protocol for identifying new ones will enable the design and testing of multiplex genetic circuits built from orthogonal LSR-RDF modules. It is anticipated that these constructs will be used in designing more complex cellular operations with applications in synthetic biology.

## DATA AVAILABILITY

The data underlying this article has been deposited in Mendeley (output of Alphafold2-multimer).

## SUPPLEMENTARY DATA

Supplementary Data are available online.

## AUTHOR CONTRIBUTIONS

Phoebe Rice and Femi Olorunniji: Conceptulization, Methodology, Writing, Funding acquisition. Heewhan Shin: Methodology, Software, Investigation. Tania Peña Reyes, Alyssa Retiz, Ying Pigli, Oluwateniola Taiwo-Aiyerin, Abdulrazak Alsaleh, Alexandria Holland, Adebayo Bello: investigation.

## ACKNOWLEDGEMENTS FUNDING

This work was supported by the National Science Foundation and UK Research and Innovation [collaborative grant NSF/BIO 2107527 and UKRI/BBSRC BB/X012085/1 to P.A.R. and F.J.O]. Funding for open access charge: National Science Foundation.

## CONFLICT OF INTEREST

none

## TABLE AND FIGURES LEGENDS

**Supplementary Figure 1. AlphaFold2 predictions for positive controls.**

a. PAE plots for positive controls. Predicted alignment error (PAE) plots from alphafold-multimer for the 5 previously known LSI – RDF pairs shown in Figure 2. For these predictions, the entire LSI sequence (rather than only the DNA binding domains) was used. See Supplementary Figure 1b for a guide to interpreting these plots.
b. Guide to interpreting PAE (predicted alignment error) plots. The PAE plot for the top model for the complex of full – length phiRV1 integrase (protein A) and its RDF (Protein B) as an example. The x and y axes represent the linear amino acide sequences of the two proteins, and the coloring represents the predicted aligment error between all each pair of amino acids (blue low; red high, calculated in two slightly different ways on the two sides of the diagonal). The on-diagonal blue regions represent the independently folded domains of proteins. DBD2 is highlighted with a yellow box, with its 3 subsections labeled N-terminal half (“2N”), coiled coil, and C-terminal half (“2C”). Its checkerboard pattern reflects the insertion of the coiled coil flanked by flexible hingers: the blue off-diagonal boxes labeled with white asterisks show that the N- and C-terminal portions of DBD2 are predicted to be well-positioned relative to each other, while the red boxes marked with black asterisks show that the coiled coil is predcted to be flexible with respect to the rest of DBD2. Finally, the blue regions marked by yellow stars show that the RDF is predicted to interact with the body of DBD2. Note the red regions between the yellow stars show that the RDF is not predicted to interact with the coiled coil itself.

**Supplementary Figure 2. Previous mutational data supports models for LSI-RDF interactions.**

Models for phiC31, Bxb1, and SPbeta integrases with their cognate RDFs are shown. Residues in either partner whose mutation interferes with RDF function or binding are shown in magenta. Note that some mutations may affect protein stability as well as protein-protein interactions. DBD2 of the integrase is shown in yellow; the CC in tan, and the RDF in cyan. The core beta strands of DBD2 are oriented similarly in all 3 pictures. (18, 22, 23)

**Supplementary Figure 3. Virtual pulldowns for positive controls**

a. Virtual Pulldown output for A118. Top: a plot similar to that in Figure 3, but with “gene name . model #” added on the horizontal axes. pTM (red) and ipTM (blue) for each of 5 models predicted for the complex of each element-encoded protein with the A118 integrase. The highest-ranking model for the known RDF is marked with a yellow star. Bottom: PAE plots for the 5 models with the known RDF. For this prediction, the last 400 amino acids of the protein were used as bait, which ensured inclusion of both DNA binding domains.
b. Virtual Pulldown output for Bxb1. See legend to Supplementary Figure 3a for detail.
c. Virtual Pulldown output for phiC31. See legend to Supplementary Figure 3a for detail.
d. Virtual Pulldown output for phiRv1. See legend to Supplementary Figure 3a for detail.
e. Virtual Pulldown output for SPbeta. See legend to Supplementary Figure 3a for detail.

**Supplementary figure 4. Orthogonality in LSR-RDF system.**

a. Currently known LSR and RDF pairs are mix to test for orthogonality. ipTM scores (0 to 1) of the highest ranked model were used to generate a heatmap. Higher the ipTM score (closre to 1) indicates the higher confidence in predicted interface between LSI and RDF models. The two darkest off-diagonal boxes can be seen to be false positives by examining the PAE plots in Supplementary figures 4c (for phiRv1 int with Bxb1’s RDF) and 4f (for A118 int with C31’s RDF) – the non-cognate RDF is not predicted to interact with DBD2 of the integrase in question. Supplementary figure 1b provides a guide to interpreting PAE plots for these predictions.
b. Predicted alignment error (PAE) plots for the SPbeta integrase with its own and 4 other RDFs.
c. Predicted alignment error (PAE) plots for the phiRv1 integrase with its own and 4 other RDFs.
d. Predicted alignment error (PAE) plots for the phiC31 integrase with its own and 4 other RDFs.
e. Predicted alignment error (PAE) plots for the Bxb1 integrase with its own and 4 other RDFs.
f. Predicted alignment error (PAE) plots for the A118 integrase with its own and 4 other RDFs.

**Supplementary Figure 5. PAE plots for the 6 integrase – RDF complexes shown in Figure 4 and experimentally tested.**

**Supplementary figure 6. Predicted orthogonality in LSR-RDF system.** LSR and RDF pairs are mixed to test for orthogonality. ipTM scores (0 to 1) of the highest ranked model were used to generate a heatmap. Higher the ipTM score (closer to 1) indicates the higher confidence in predicted interface between LSI and RDF models.

The two darkest off-diagonal boxes can be seen to be false positives by examining the PAE plots the supplementary data.

**Supplementary figure 7. In vivo recombination reactions of Bt24, Cb16, and Int30 and Int10 integrases and their fusions with putative RDFs identified through virtual pulldown.** See Main Figure 6 for details.

**Supplementary Figure 8: *In vitro* recombination reactions of Int30 integrase and its fusion with the RDF.** (A) Schematic illustration of the *in vitro* recombination (inversion) assay. LSI-catalysed recombination (inversion) of *attP* and *attB* sites in pInt30-invPB gives rise to *attR* and *attL* sites in the product plasmid (pInt30-invRL), and vice versa. (B) Reactions were carried out for 2 hours as described in Materials and Methods. Integrase and integrase-RDF fusions were used at two final concentrations of 0.4, 0.8, and 1.6 µM. Reaction products were digested with the restriction endonuclease, XhoI, prior to 1.2% agarose gel electrophoresis. As illustrated in **A**, digestion of the plasmids with XhoI gives different restriction patterns for recombined and non-recombined DNA. The bands on the gel are labeled *nr* (non-recombinant, i.e. substrate), *rec* (recombination product).

**Supplementary Table 1: LSRs tested and their predicted RDFs.** This table contains the LSRs used in virtual pulldowns (10) (31), their amino acid sequences, putative RDFs identified for them, and their locus tags and amino acid sequences.

## Notes

### Competing Interest Statement

The authors have declared no competing interest.

