## Supplementary Figures for "Identification of cognate recombination directionality factors for large serine recombinases by virtual pulldown"

List of supplementary figures


1. PAE plots for 5 positive controls
   1a PAE plots
   1b Guide to interpreting PAE plots

2. Previous mutational data supports models for LSI-RDF interactions

3. Virtual pulldowns for 5 positive controls
   3a for A118
   3b for Bxb1
   3c for phiC31
   3d for phiRv1
   3e SPbeta

4. Predicted orthogonality for the 5 positive control pairs
   4a Plot of all vs. all ipTM scores
   4b PAE plots for SPbeta int + 5 RDFs
   4c for phiRv1
   4d for phiC31
   4e for Bxb1
   4f for A118

5. PAE plots for experimental pulldowns for the 6 RDFs experimentally tested

6. Predicted orthogonality for selected predicted RDFs

*7.* *in vivo* data for Bt24, Cb16, Int10 and Int30

*8.* *in vitro* data for Int30

### Supplementary Fig. 1a (Related to Fig. 2)

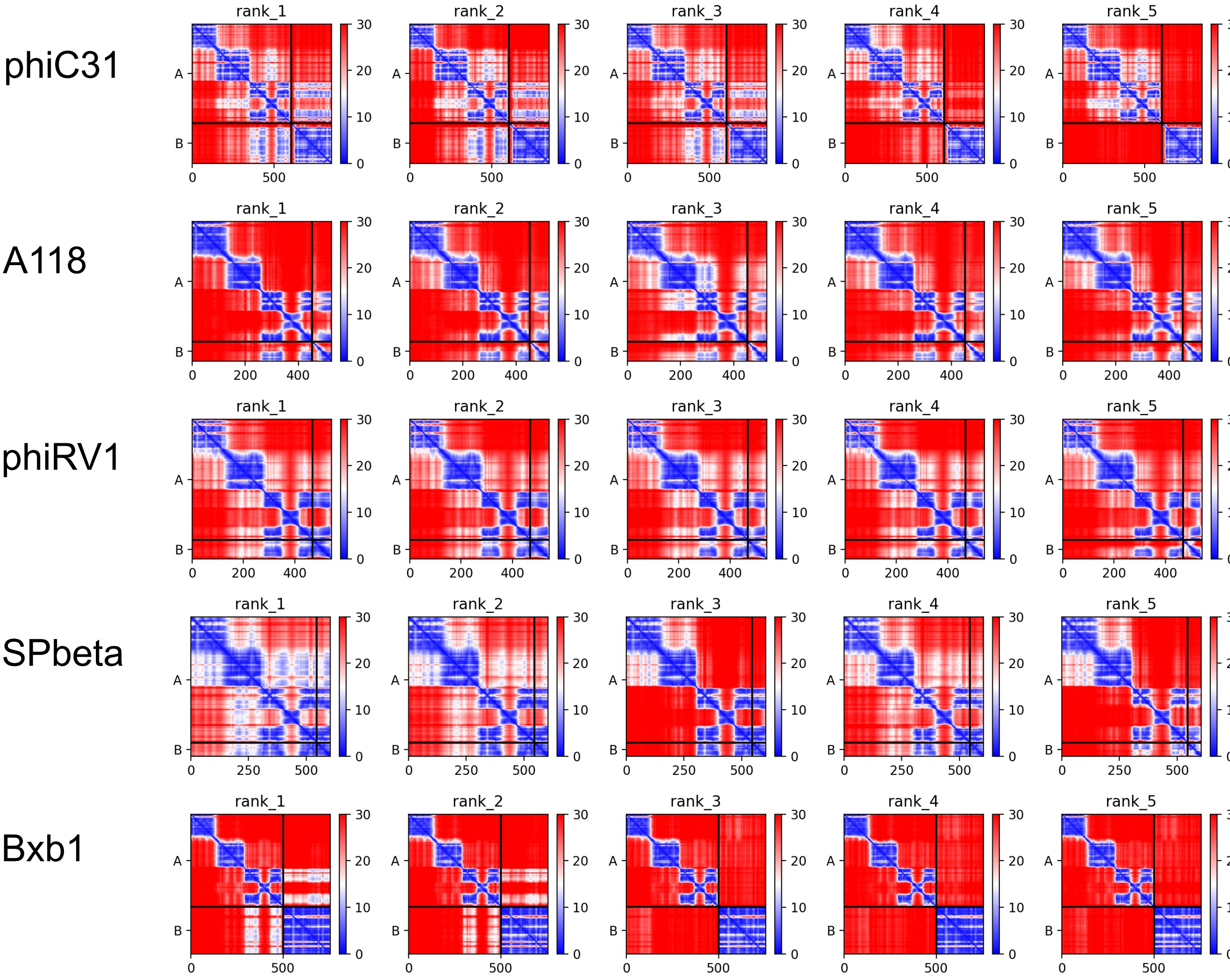


Supplementary Figure 1a. PAE plots for positive controls.
Predicted alignment error (PAE) plots from alphafold-multimer for the 5 previously known LSI – RDF pairs shown in Figure 2. For these predictions, the entire LSI sequence (rather than only the DNA binding domains) was used.
See Supplementary Figure 1b for a guide to interpreting these plots.

**Supplementary Fig. 1b (Related to Fig. 2)**

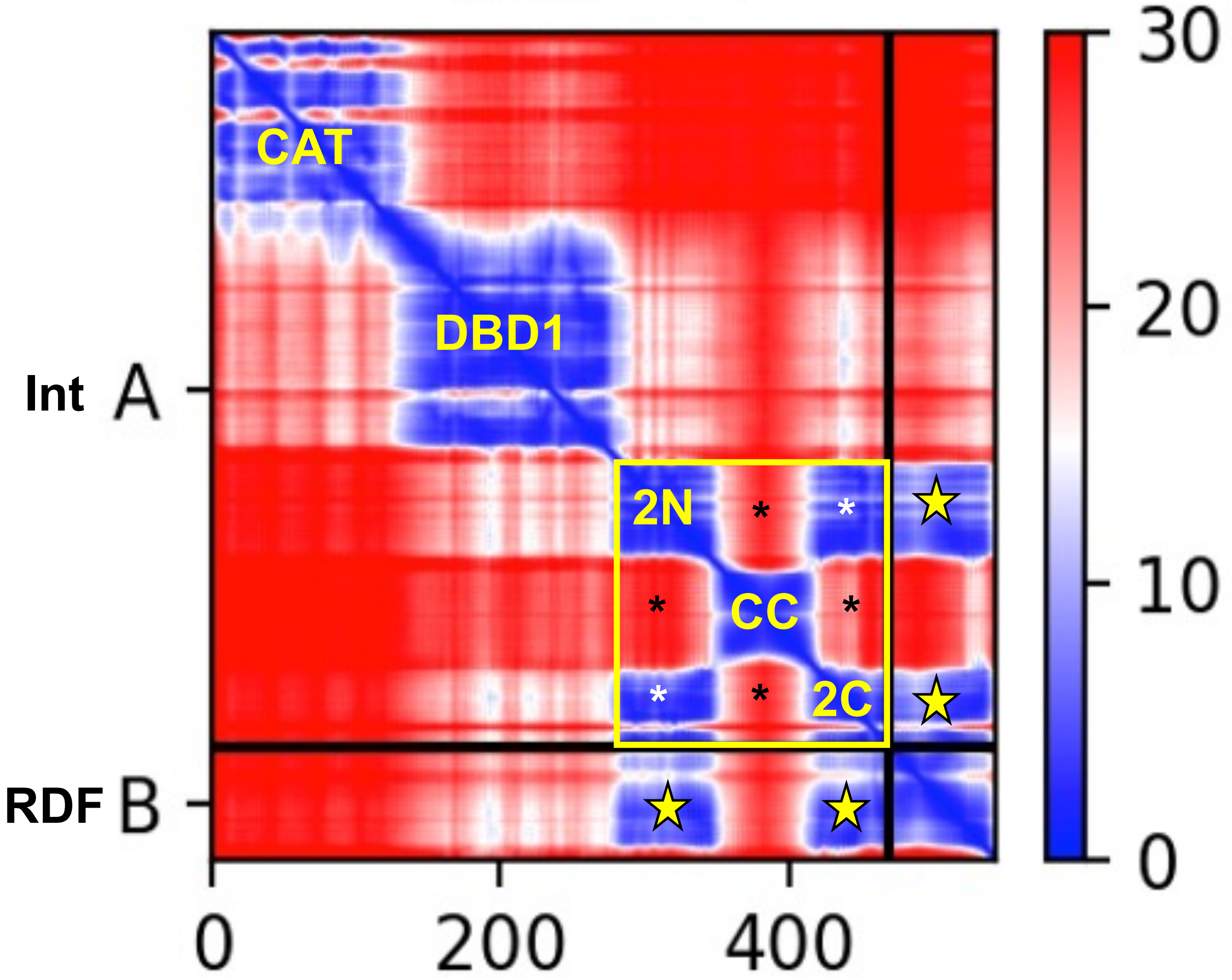


Supplementary Figure 1b. Guide to interpreting PAE (predicted alignment error) plots.

The PAE plot for the top model for the complex of full – length phiRv1 integrase (protein A) and its RDF (Protein B) as an example. The x and y axes represent the linear amino acide sequences of the two proteins, and the coloring represents the predicted aligment error between all each pair of amino acids (blue low; red high, calculated in two slightly different ways on the two sides of the diagonal).

The on-diagonal blue regions represent the independently folded domains of proteins. DBD2 is highlighted with a yellow box, with its 3 subsections labeled N-terminal half ("2N"), coiled coil, and C-terminal half ("2C"). Its checkerboard pattern reflects the insertion of the coiled coil flanked by flexible hingers: the blue off-diagonal boxes labeled with white asterisks show that the N- and C-terminal portions of DBD2 are predicted to be well-positioned relative to each other, while the red boxes marked with black asterisks show that the coiled coil is predcted to be flexible with respect to the rest of DBD2.

Finally, the blue regions marked by yellow stars show that the RDF is predicted to interact with the body of DBD2. Note the red regions between the yellow stars show that the RDF is not predicted to interact with the coiled coil itself.

### Supplementary Fig. 2 (Related to Fig. 2)

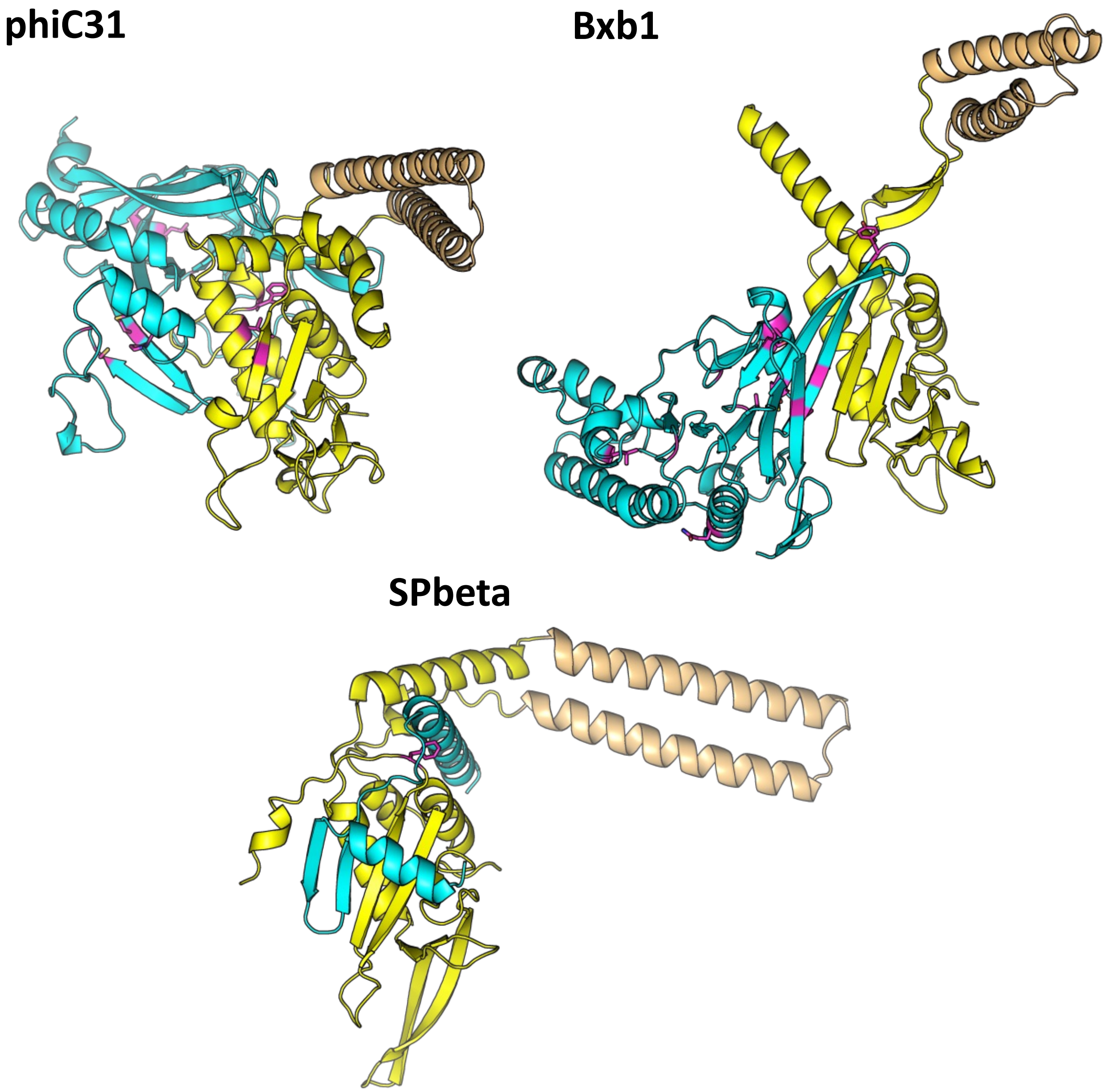


Supplementary Figure 2. Previous mutational data supports models for LSI-RDF interactions. Models for phiC31, Bxb1, and SPbeta integrases with their cognate RDFs are shown. Residues in either partner whose mutation interferes with RDF function or binding are shown in magenta. Note that some mutations may affect protein stability as well as protein-protein interactions. DBD2 of the integrase is shown in yellow; the CC in tan, and the RDF in cyan. The core beta strands of DBD2 are oriented similarly in all 3 pictures. (18, 22, 23)

### Supplementary Fig. 3a

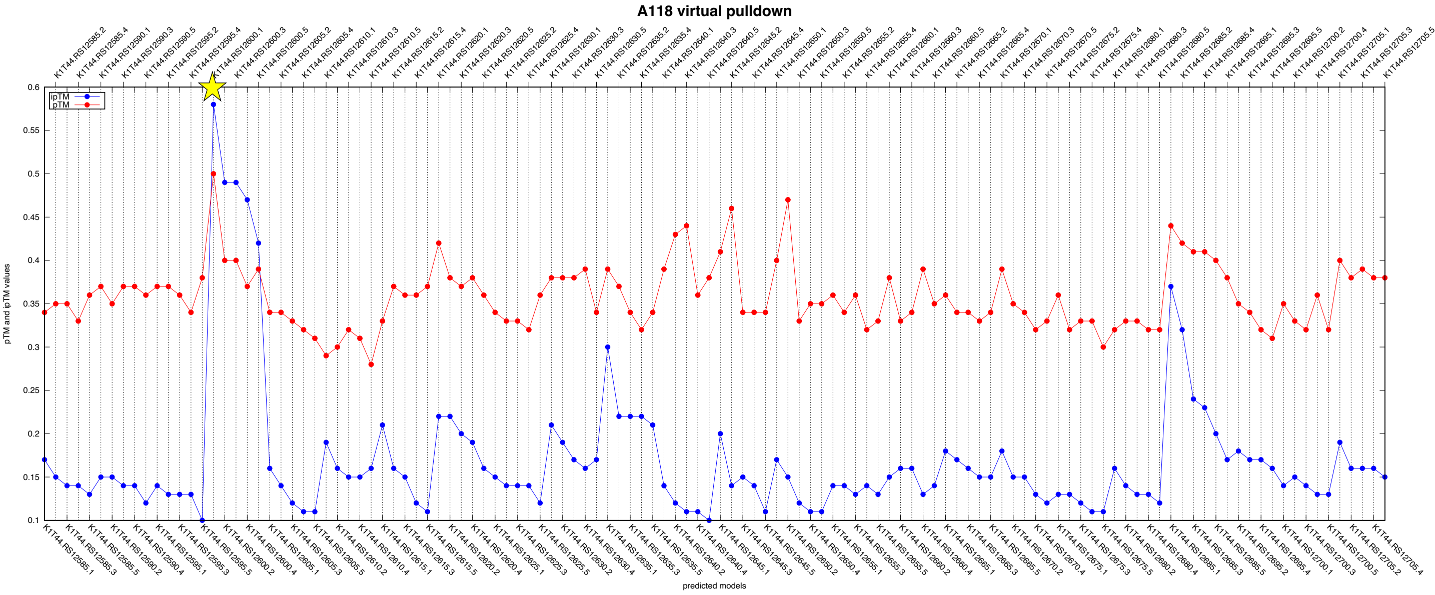


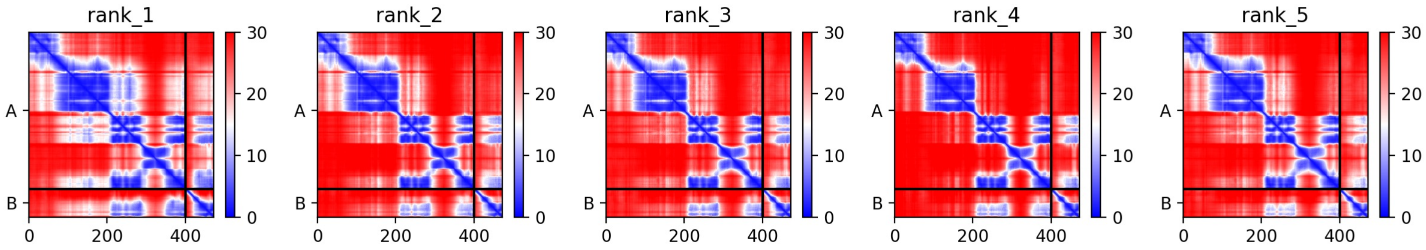


### Supplementary Figure 3. Virtual pulldowns for positive controls

Supplementary Figure 3a. Virtual Pulldown output for A118.
Top: a plot similar to that in Figure 3, but with “gene name . model #” added on the horizontal axes. pTM (red) and ipTM (blue) for each of 5 models predicted for the complex of each element-encoded protein with the A118 integrase. The highest-ranking model for the known RDF is marked with a yellow star.
Bottom: PAE plots for the 5 models with the known RDF. For this prediction, the last 400 amino acids of the protein were used as bait, which ensured inclusion of both DNA binding domains.

### Supplementary Fig. 3b, continued.

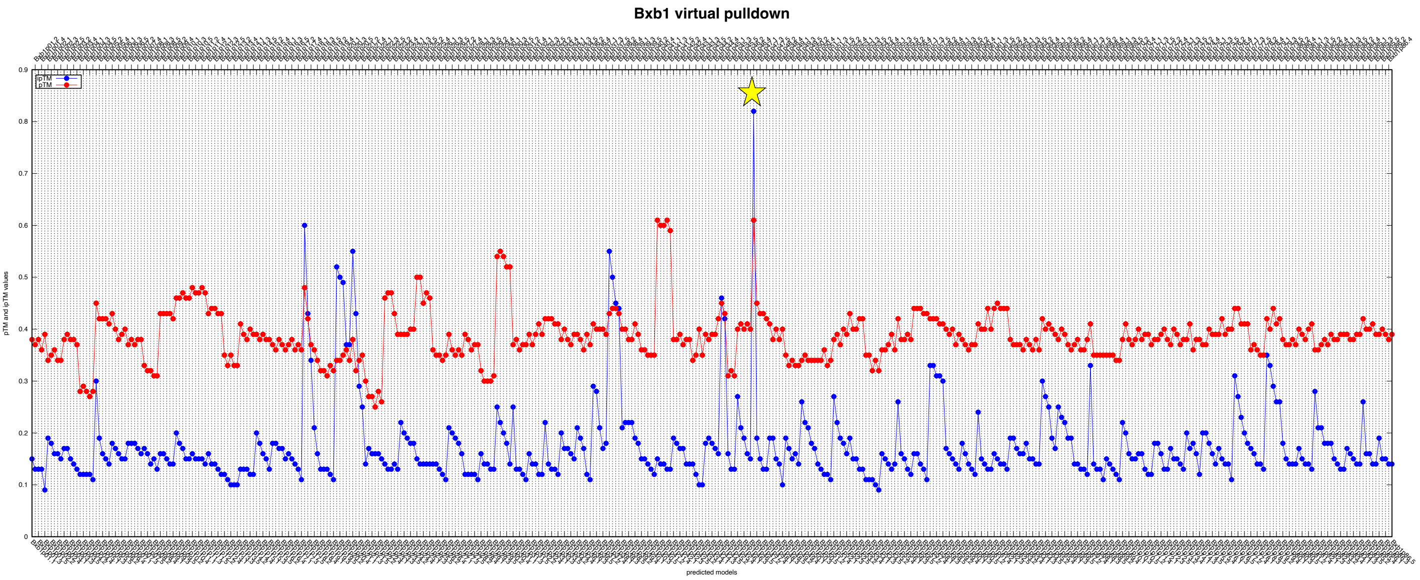


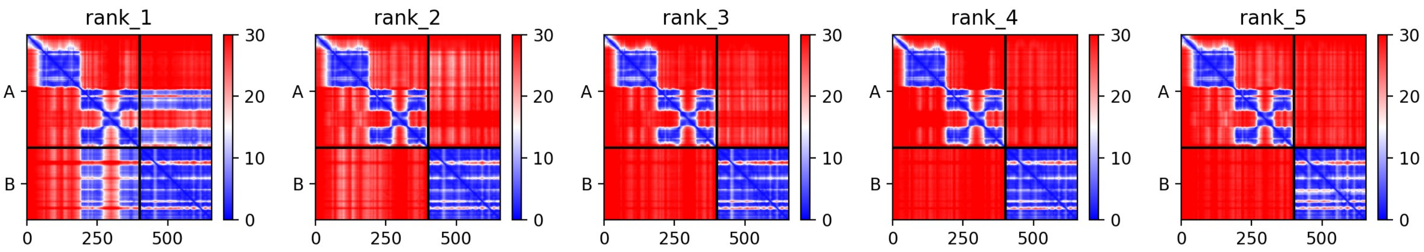


Supplementary Figure 3b. Virtual Pulldown output for Bxb1.
See legend to Supplementary Figure 3a for detail.

### Supplementary Fig. 3c, continued.

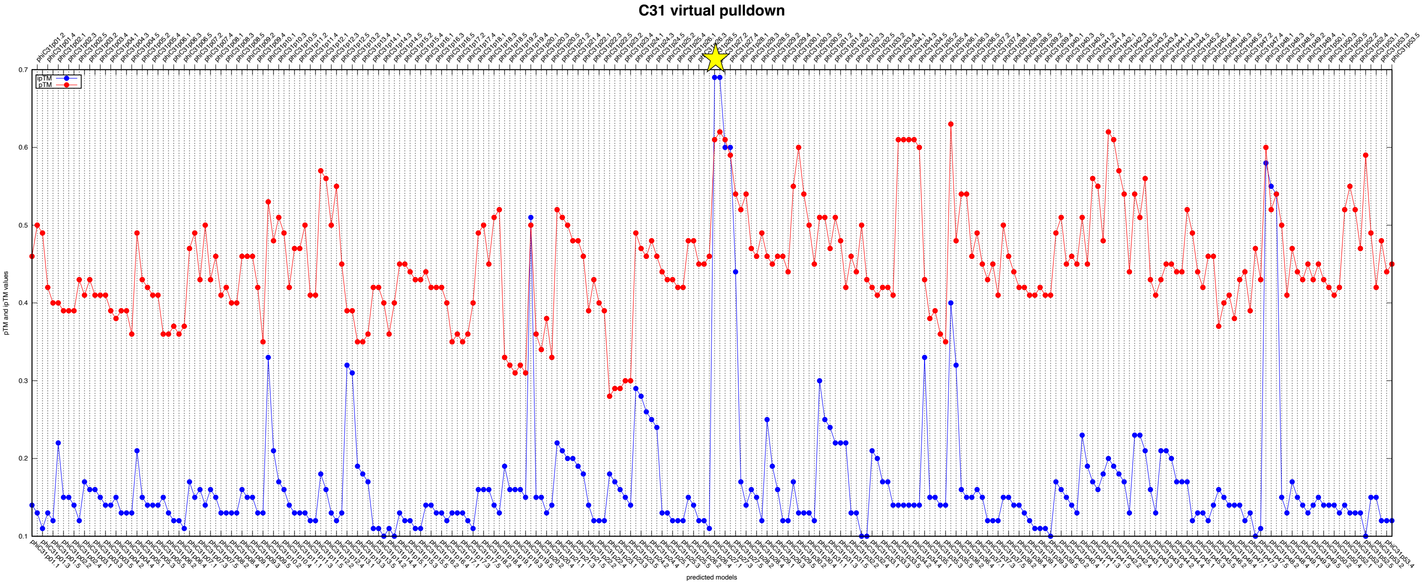


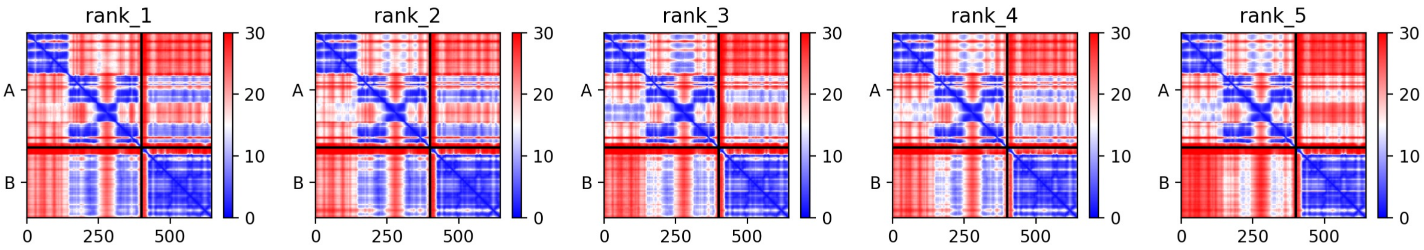


Supplementary Figure 3c. Virtual Pulldown output for phiC31.
See legend to Supplementary Figure 3a for detail.

### Supplementary Fig. 3d, continued.

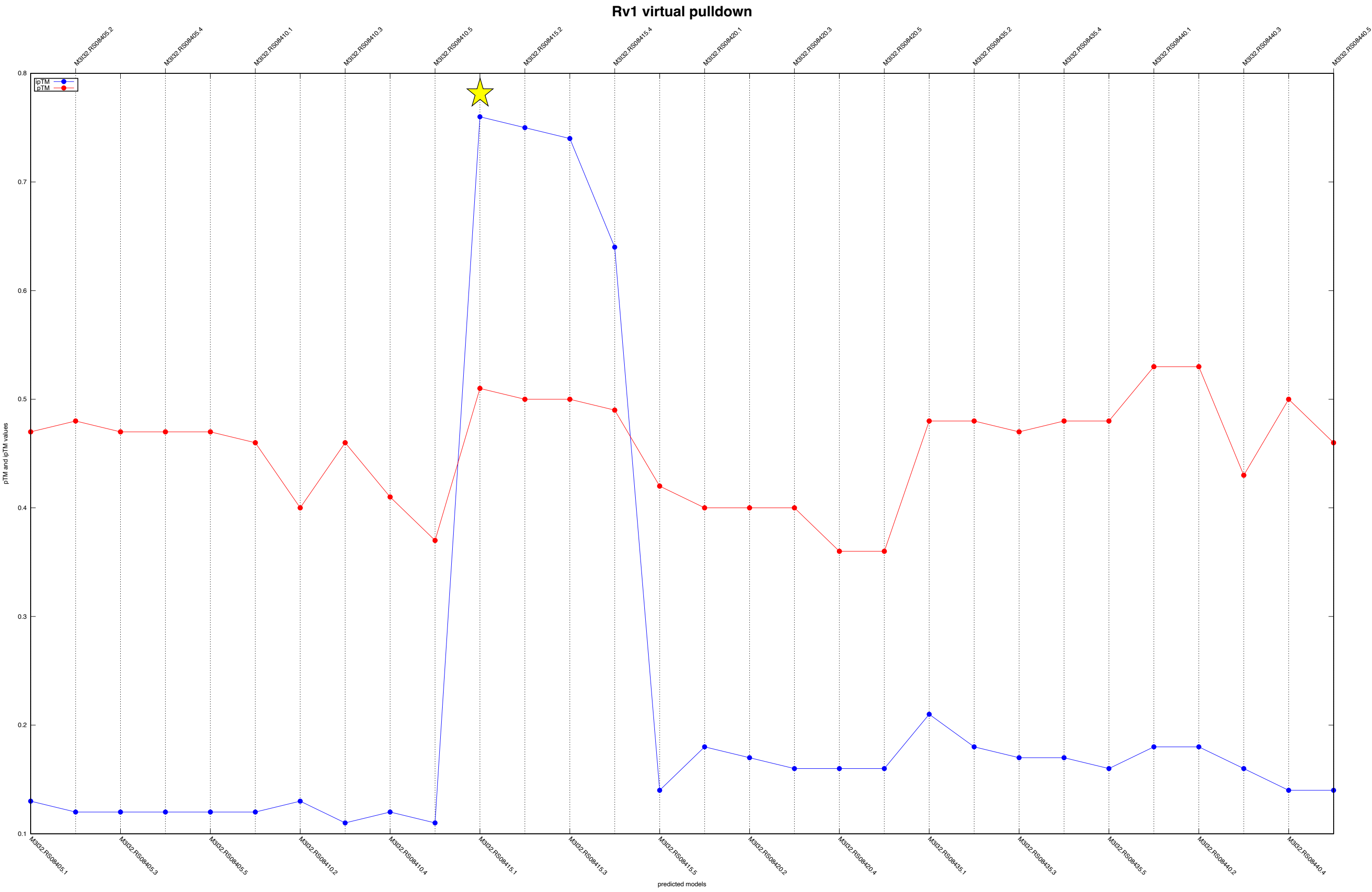


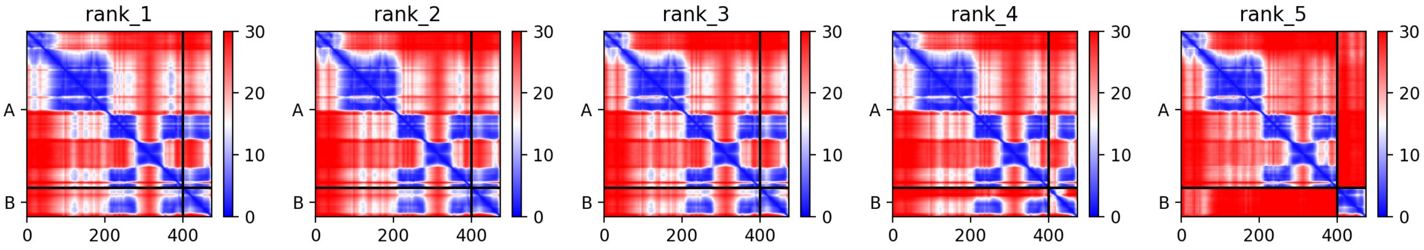


Supplementary Figure 3d. Virtual Pulldown output for phiRv1. See legend to Supplementary Figure 3a for detail.

### Supplementary Fig. 3e, continued.

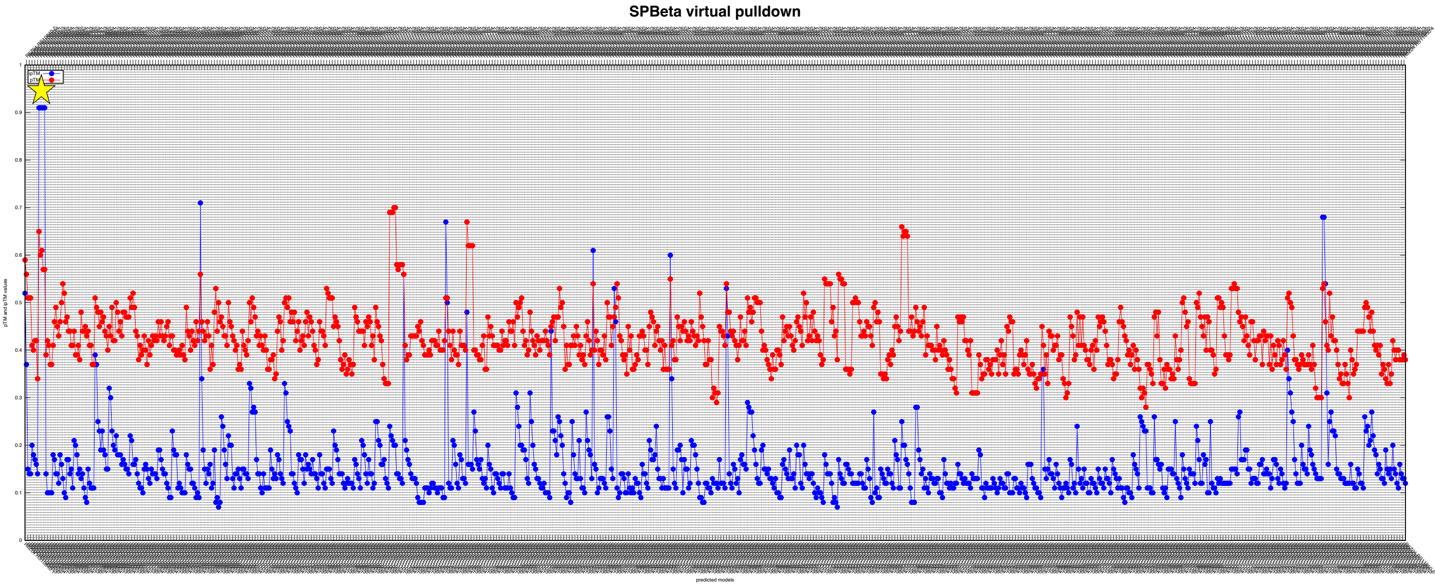


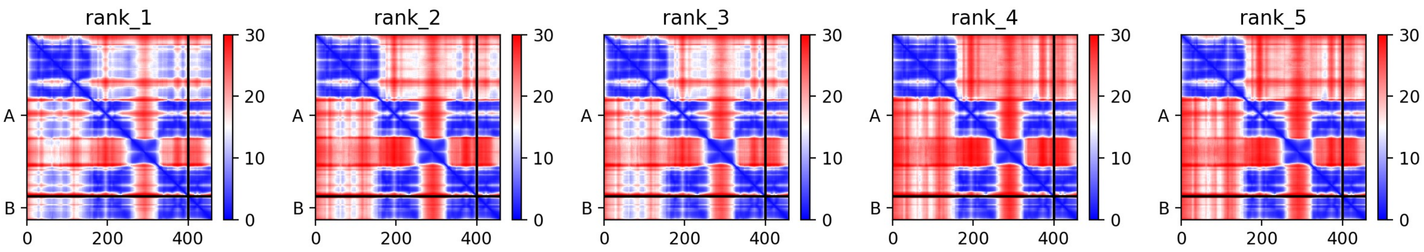


Supplementary Figure 3e. Virtual Pulldown output for SPbeta.
See legend to Supplementary Figure 3a for detail.

### Supplementary Fig. 4a

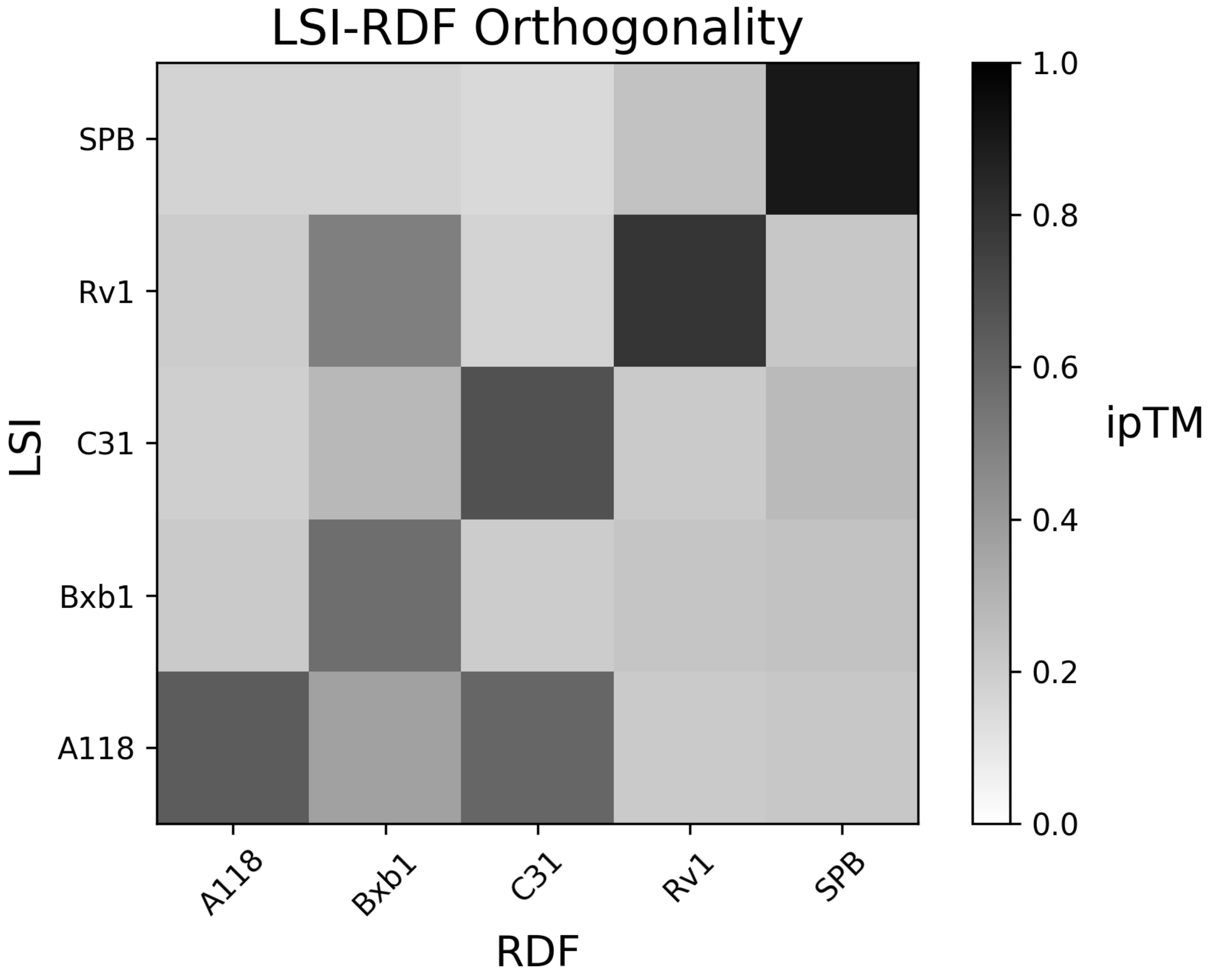


**Supplementary figure 4. Predicted orthogonality for the 5 positive control pairs**

Supplementary figure 4a. Orthogonality in LSR-RDF system. Currently known LSR and RDF pairs are mix to test for orthogonality. ipTM scores (0 to 1) of the highest ranked model were used to generate a heatmap. Higher the ipTM score (closre to 1) indicates the higher confidence in predicted interface between LSI and RDF models.

The two darkest off-diagonal boxes can be seen to be false positives by examining the PAE plots in Supplementary figures 4c (for phiRv1 int with Bxb1's RDF) and 4f (for A118 int with phiC31's RDF) – the non-cognate RDF is not predicted to interact with DBD2 of the integrase in question.  Supplementary figure 1b provides a guide to interpreting PAE plots for these predictions.

### Supplementary Fig. 4b, continued.

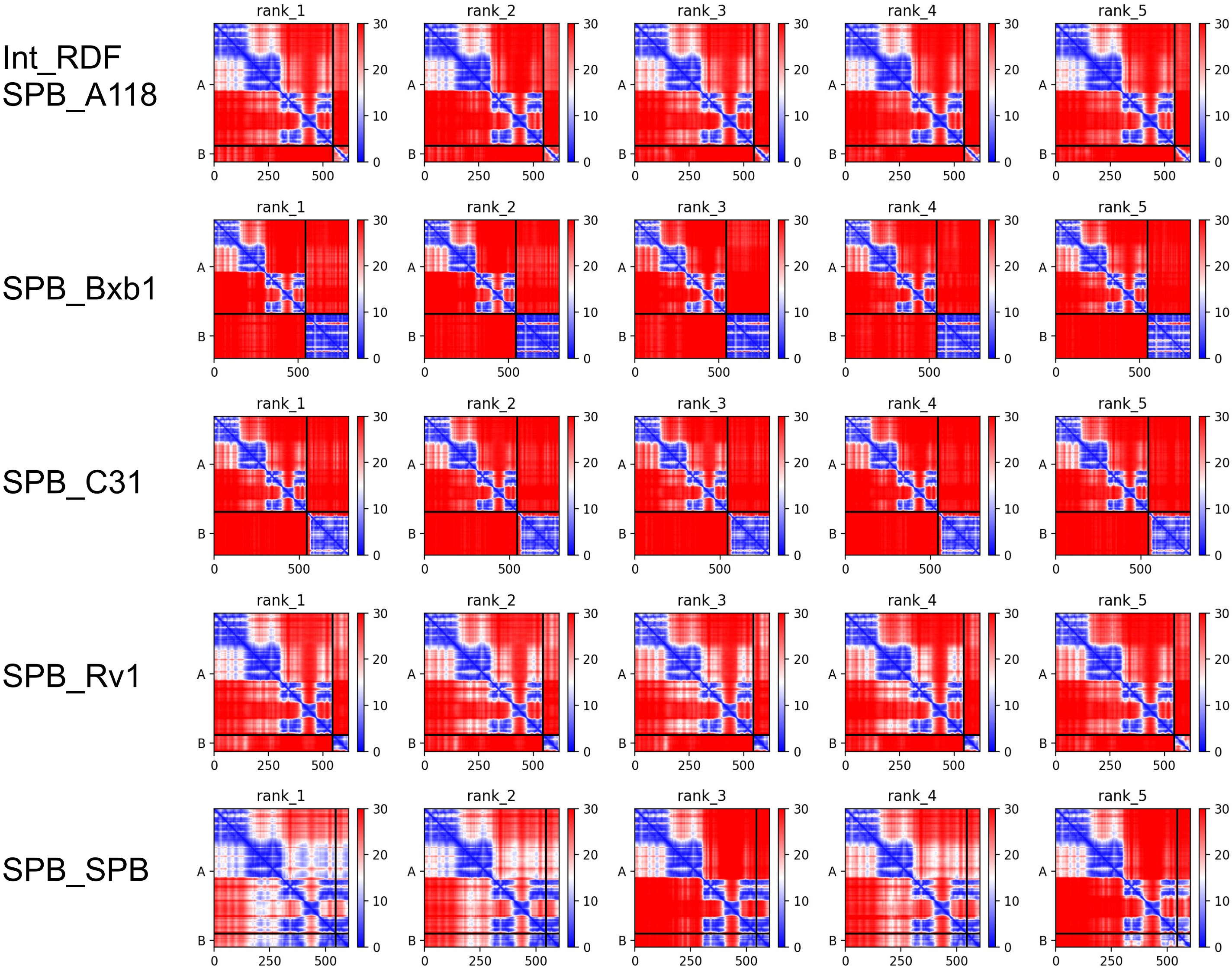


Supplementary figure 4b. Orthogonality in LSI-RDF system. Predicted alignment error (PAE) plots for the phage SPbeta integrase with its own and 4 other RDFs.

### Supplementary Fig. 4c, continued.

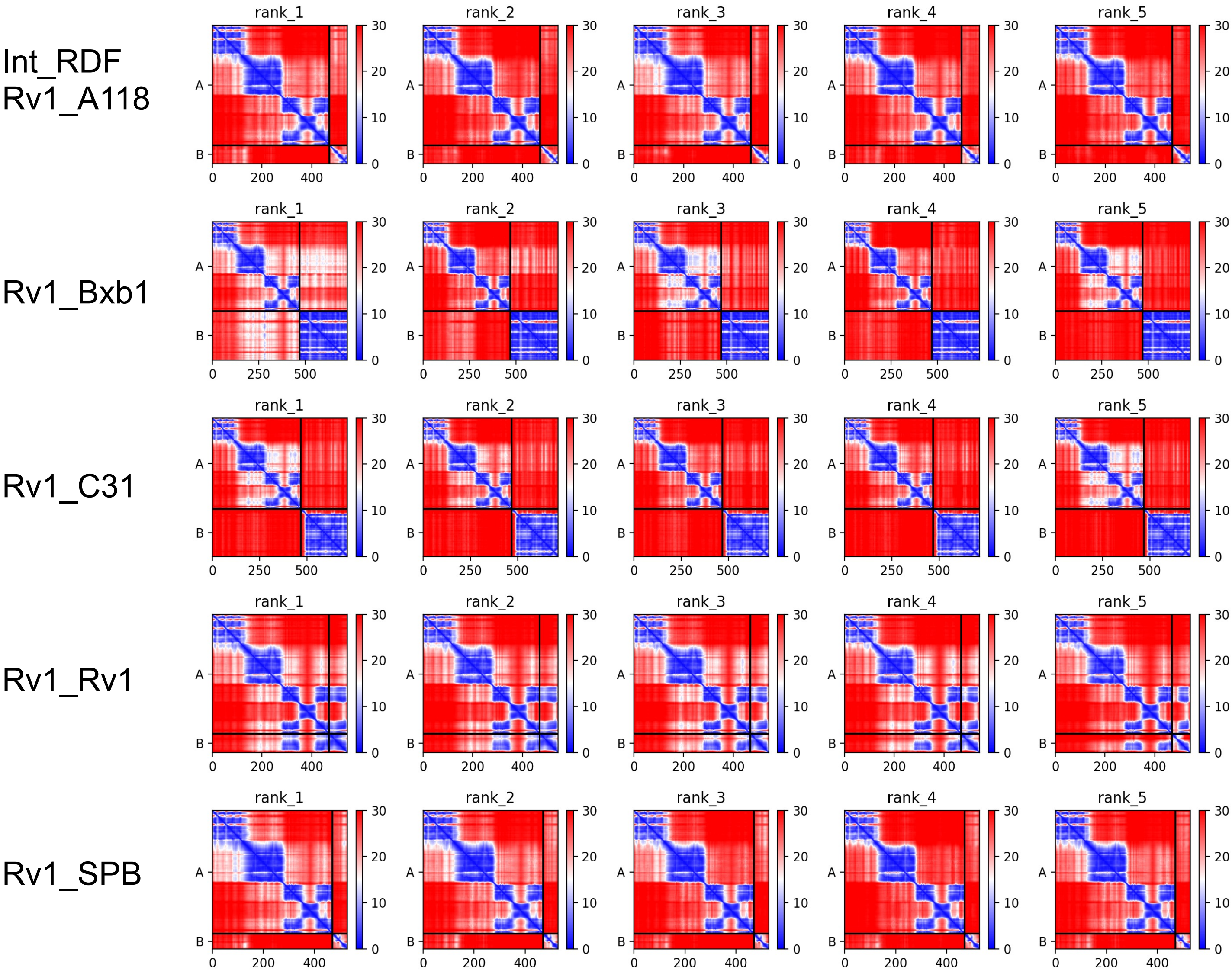


Supplementary figure 4c. Predicted alignment error (PAE) plots for mixed LSI-RDF pairs for the phiRv1 integrase.

### Supplementary Fig. 4d, continued.

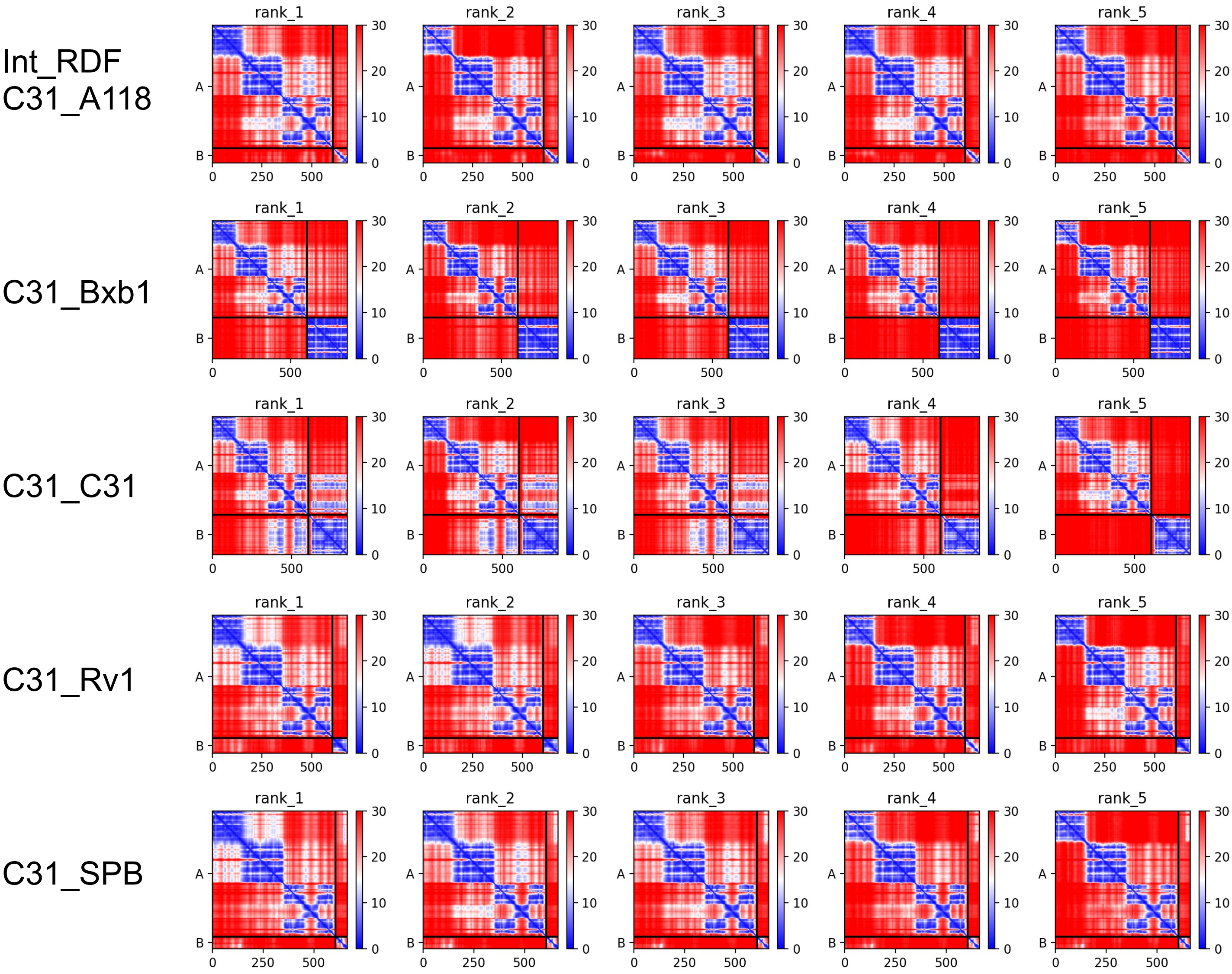


Supplementary figure 4d. Predicted alignment error (PAE) plots for mixed LSI-RDF pairs for the phiC31 integrase.

### Supplementary Fig. 4e, continued.

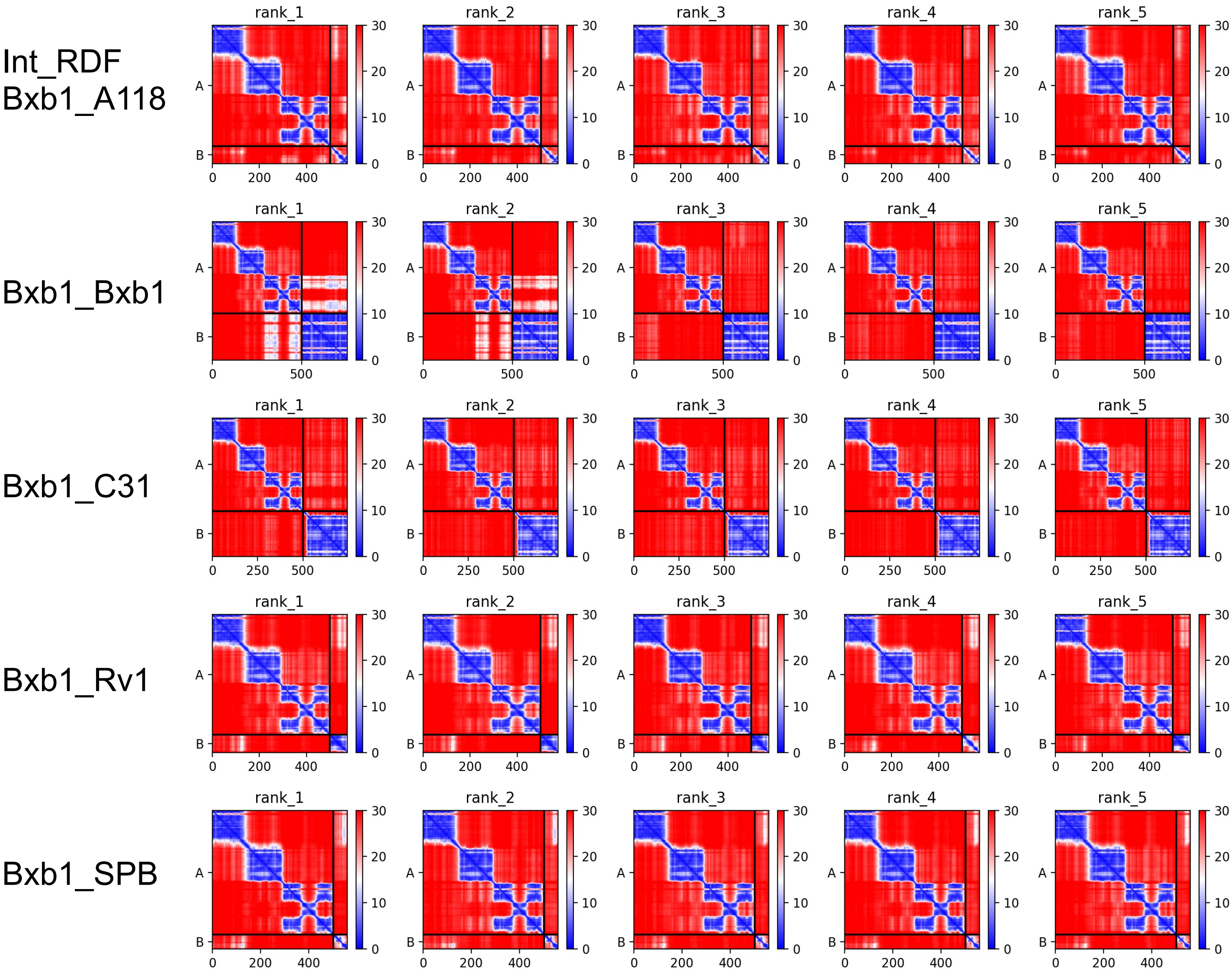


Supplementary figure 4e. Predicted alignment error (PAE) plots for mixed LSI-RDF pairs for the Bxb1 integrase.

### Supplementary Fig. 4f, continued.

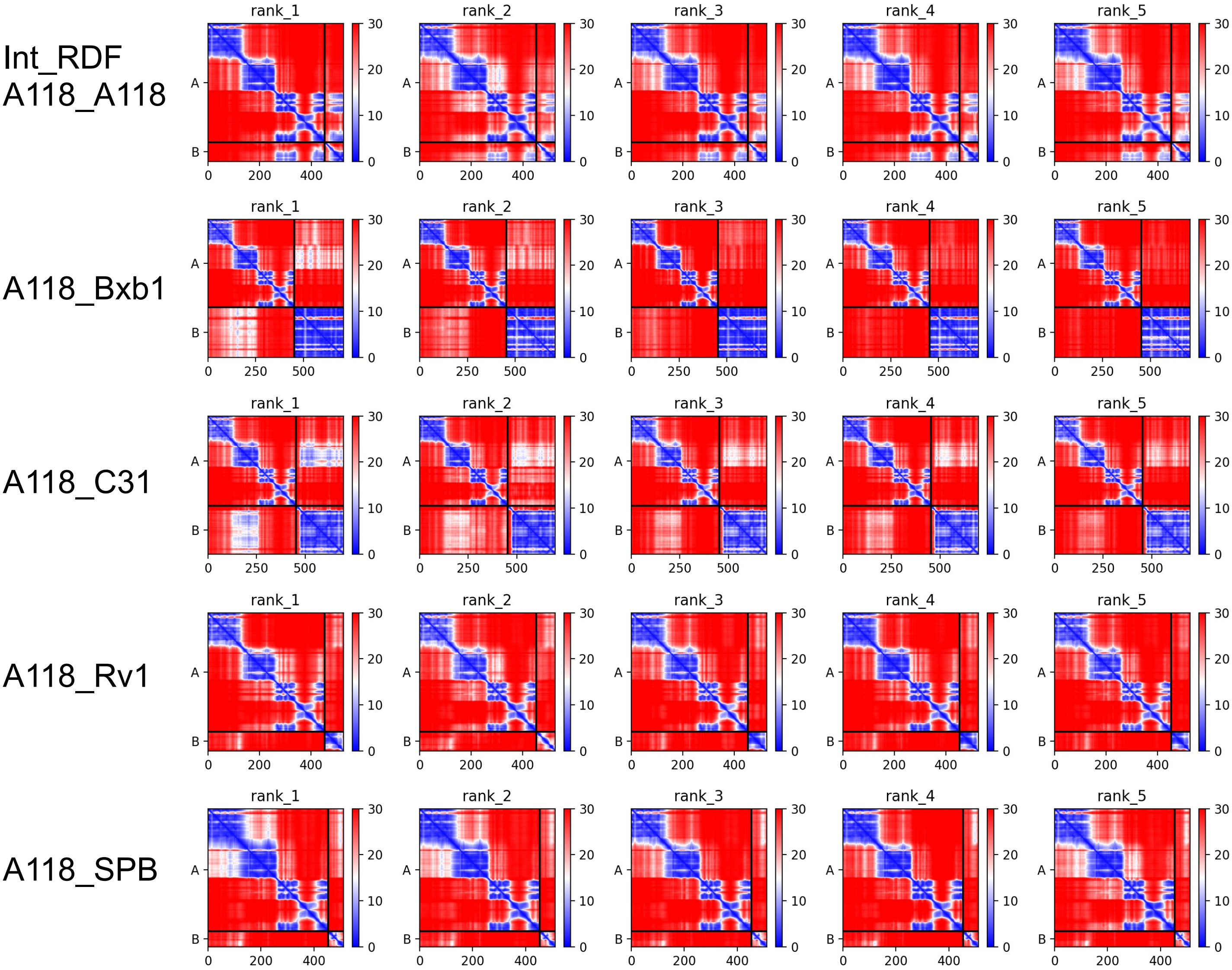


Supplementary figure 4f. Orthogonality in LSI-RDF system. Predicted alignment error (PAE) plots for mixed LSI-RDF pairs for the A118 integrase.

### Supplementary Fig. 5

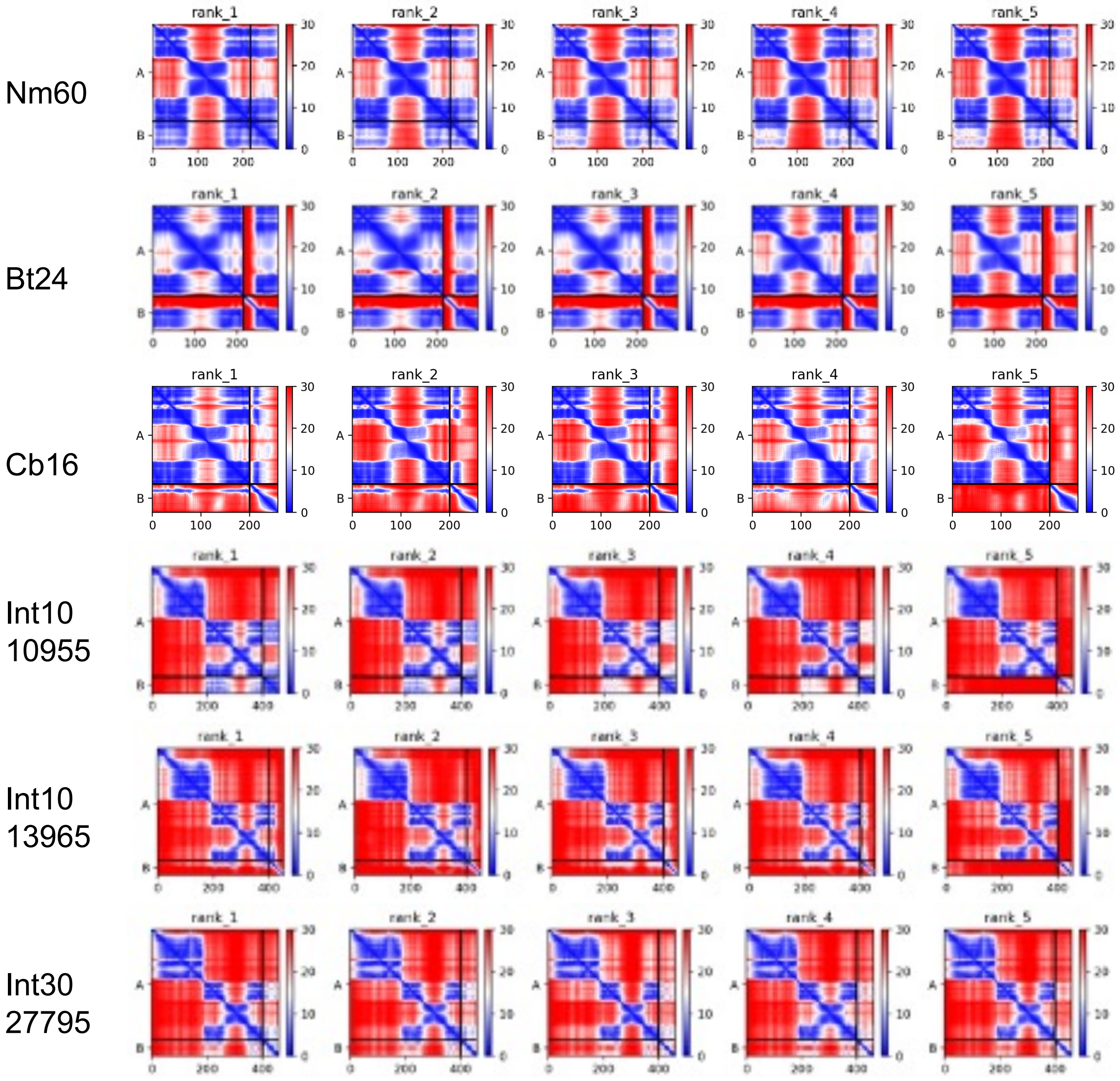


Supplementary Figure 5. PAE plots for the 6 integrase – RDF complexes shown in Figure 4 and experimentally tested.

#### Supplementary Fig. 6

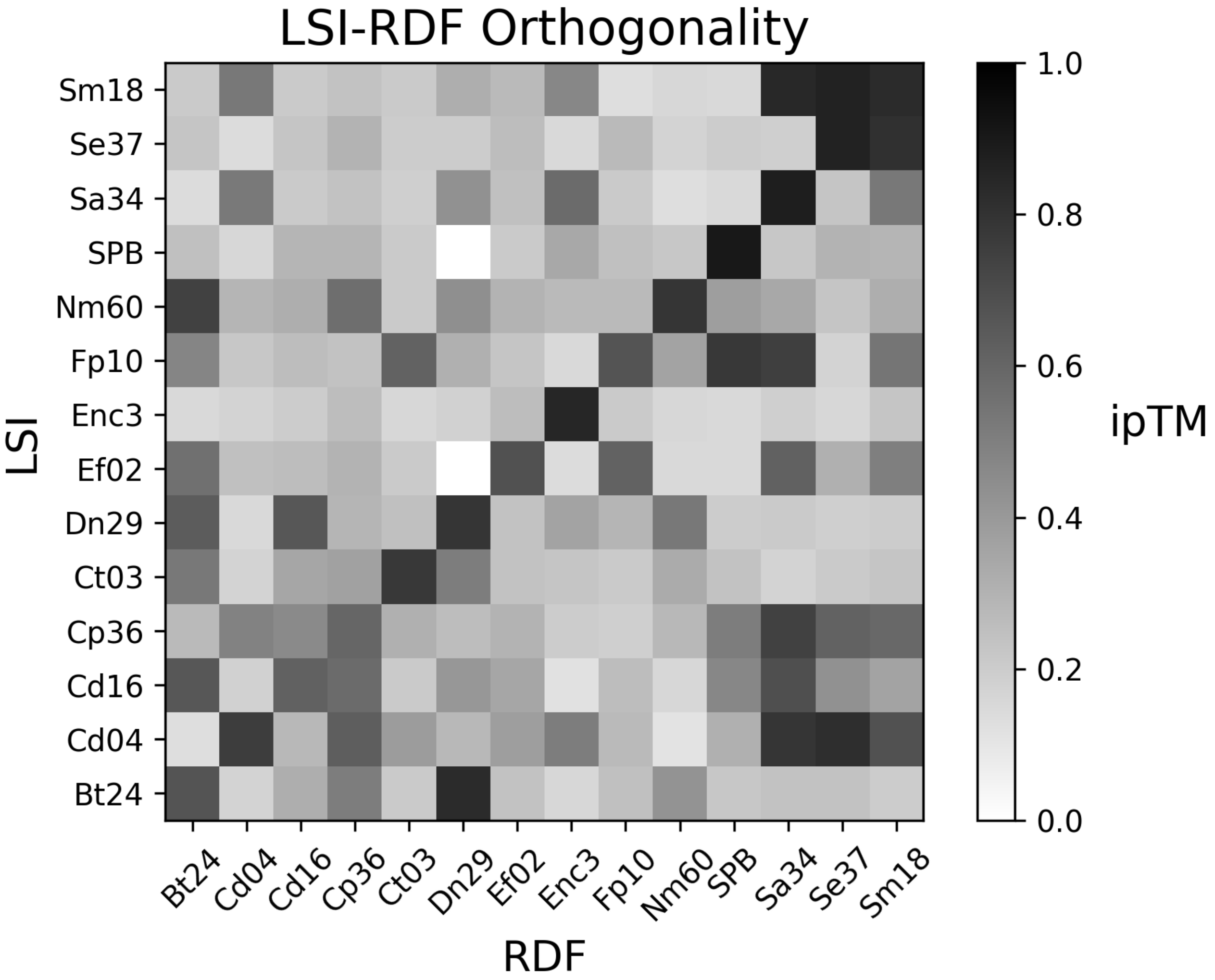


Supplementary figure 6. Predicted orthogonality in LSR-RDF system. LSR and RDF pairs are mixed to test for orthogonality. ipTM scores (0 to 1) of the highest ranked model were used to generate a heatmap. Higher the ipTM score (closer to 1) indicates the higher confidence in predicted interface between LSI and RDF models.

The two darkest off-diagonal boxes can be seen to be false positives by examining the PAE plots the supplementary data.

### Supplementary Fig. 7

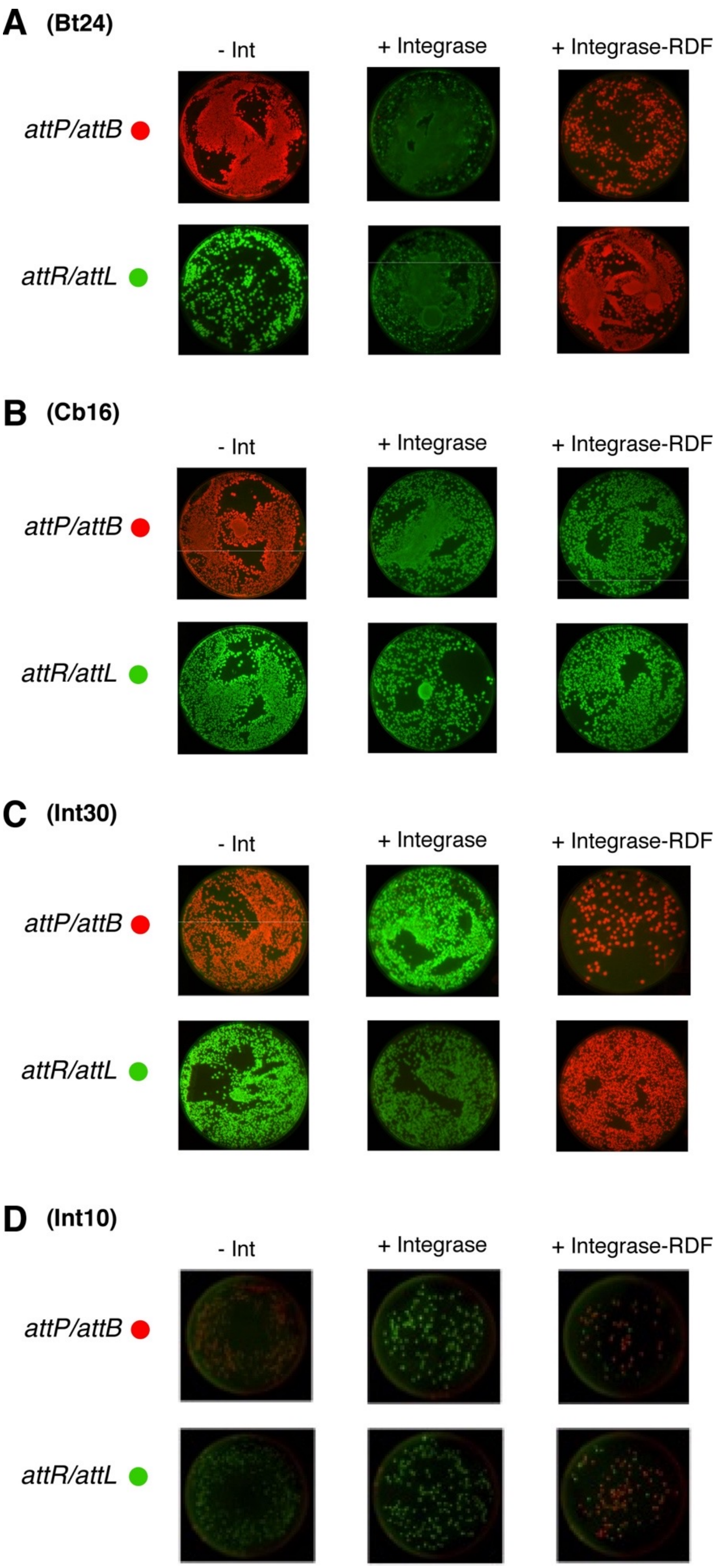


**Supplementary figure 7**. *In vivo* recombination reactions of Bt24, Cb16, and Int30 and Int10 integrases and their fusions with putative RDFs identified through virtual pulldown. See Main Figure 6 for details.

### Supplementary Fig. 8

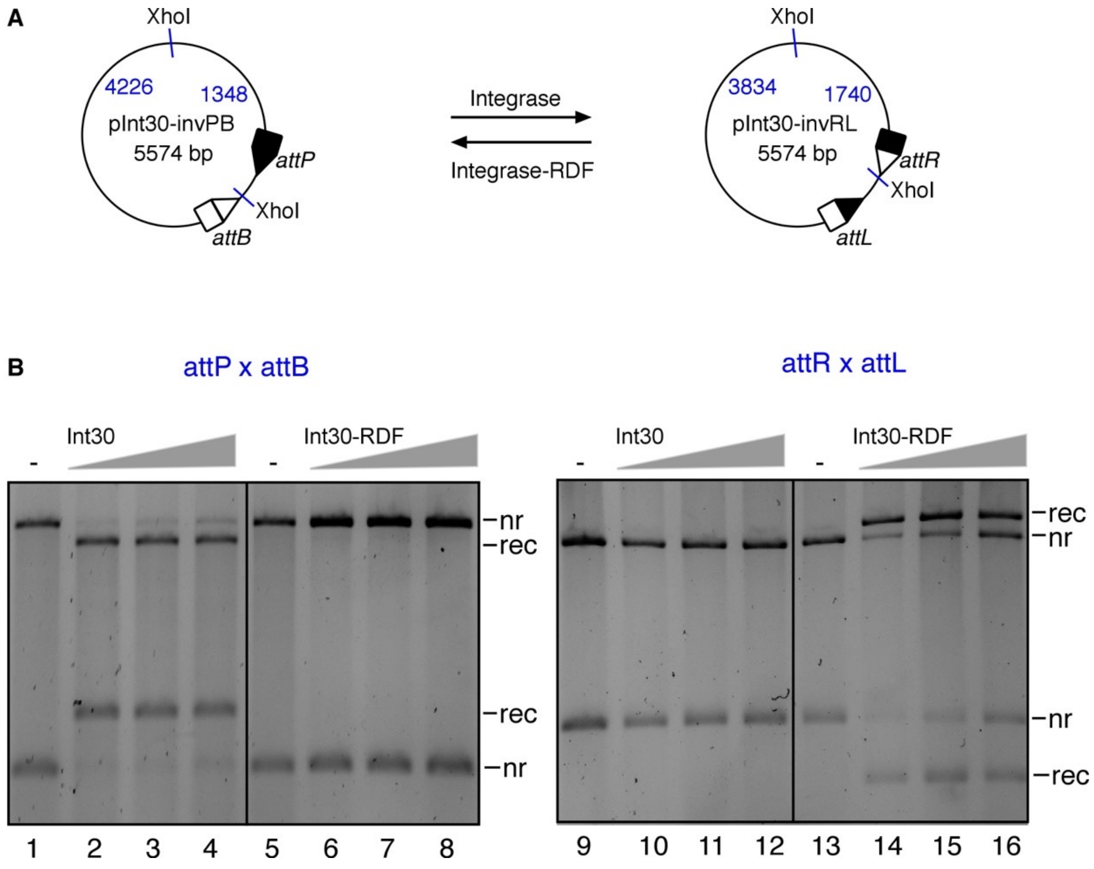


**Supplementary Figure 8: *In vitro* recombination reactions of Int30 integrase and its fusion with the RDF.** (A) Schematic illustration of the *in vitro* recombination (inversion) assay. LSI-catalysed recombination (inversion) of *attP* and *attB* sites in pInt30-invPB gives rise to *attR* and *attL* sites in the product plasmid (pInt30-invRL), and vice versa. (B) Reactions were carried out for 2 hours as described in Materials and Methods. Integrase and integrase-RDF fusions were used at two final concentrations of 0.4, 0.8, and 1.6 μM. Reaction products were digested with the restriction endonuclease, XhoI, prior to 1.2% agarose gel electrophoresis. As illustrated in **A**, digestion of the plasmids with XhoI gives different restriction patterns for recombined and non-recombined DNA. The bands on the gel are labeled *nr* (non-recombinant, i.e. substrate), *rec* (recombination product).
