## Supplementary Table 1 for "Identification of cognate recombination directionality factors for large serine recombinases by virtual pulldown"

**Supplementary Table 1: LSRs tested and their predicted RDFs.** This table contains the LSRs used in virtual pulldowns (10) (31), their amino acid sequences, putative RDFs identified for them, and their locus tags and amino acid sequences. "Yes" in the "DBD2_only" column signifies that the RDF was found using DBD2 only as bait; "no" signifies that both DNA binding domains were included in the bait.

10. Durrant,M.G., Fanton,A., Tycko,J., Hinks,M., Chandrasekaran,S.S., Perry,N.T., Schaepe,J., Du,P.P., Lotfy,P., Bassik,M.C., *et al.* (2023) Systematic discovery of recombinases for efficient integration of large DNA sequences into the human genome. *Nat Biotechnol*, **41**, 488–499.

31. Yang,L., Nielsen,A.A.K., Fernandez-Rodriguez,J., McClune,C.J., Laub,M.T., Lu,T.K. and Voigt,C.A. (2014) Permanent genetic memory with >1-byte capacity. *Nat Methods*, **11**, 1261–1266.

| lsr_id | lsr_seq | rdf_id | rdf_seq | DBD2_only |
|---|---|---|---|---|
| Sh25 | MKVAIYTRVS | ADG59_RS11785 | MLKKLKIALLIVILAEEIRS | yes |
| Bm99 | MAKKPKAKV | BMULJ_RS23410 | MTKKQSKHAEPTTDMQ | no |
| Nm60 | MSRPTGLTID | CHR53_RS06930 | MKEFKFGNATVIIHSPLV | yes |
| Cc91 | MSRRRALAP | M768_RS14940 | MMDTVVMPREEALTRA | yes |
| Bt24 | MKTAIYLRKS | FS868_RS17680 | MPSTNMAVPTESSHKH | yes |
| Bu30 | MAAKSRVYS | WJ32_RS33200 | MTEQSNFEISAVSYKGL | no |
| Ma05 | MRVLGRVRL | BUV17_RS13710 | MTRRVAIISDIQWPYHD | yes |
| Rh64 | MESASPGLR | REQ_RS23770 | MKSMLPPPTDNFDRRD | yes |
| Cb16 | MLRIAIYSRKS | ADU81_RS15025 | MAKRGRKSTAKKPKVIV | yes |
| uCb4 | MNAIYARQS | KH198_02355 | MIRIENLTPQLTAEECME | no |
| Ec03 | MRSVITYLRY | EL023_RS13870 | MKNKHTLTRITKTAFDEL | yes |
| Ef01 | MRTGLYVRV | SSG_RS18495 | MDLYAIEIRKYNKKTKRY | yes |
| Ef02 | MKRVALYMR | OMC_RS0106405 | MSNKPKITLNGTPNLQR | yes |
| Kp01 | MLRPICYERV | AD96_RS20885 | MTVSKRGRVPQTIKEAI | yes |
| Kp01 | MLRPICYERV | AD96_RS28860 | MDTVEELNGTYFYDGS | yes |
| Kp04 | MPKAISYIRFS | D2618_RS04945 | MSIPMKGLSKRGSYNHL | yes |
| Kp05 | MKKIIPYGYLF | KPLM21_RS24895 | MDVRGFLFFSVLNFLN | yes |
| Pa01 | MPSAFSYVRI | NS72_RS09065 | MSVLISRKHWDSLLLEIE | yes |
| Pa03 | MPTAYAYIRY | IPC1182_RS19485 | MSEDNRKFDKYVGEIND | no |
| Pf13 | MPKAISYIRFS | SRM1_RS29915 | MRVQLNERAFFQLEELN | no |
| Se37 | MNKVAIYVR | BMZ65_RS10355 | MAKKHYKIIHVMADGTE | yes |
| Ct03 | MENVCIYLRK | HMPREF1092_RS04285 | MMEADVTIIEPNNTAEE | yes |
| Cd31 | MPRIRKDKM | A4Z18_RS04765 | MITKLQLQQMRNVDITC | yes |
| Ps40 | MIAAIYSRKSI | UMC4401_RS04915 | MAPRKSREIKVEVVYPEI | yes |
| Ps40 | MIAAIYSRKSI | UMC4401_RS04965 | MLKKFCRCGKIIPQEISM | yes |
| Enc3 | MMKKIAVLY | FL153_RS23045 | MKQETKWTIRVFGVSFI | yes |
| Fp10 | MTENNNRV | CGS53_RS04355 | MIDKFSQVAENFPAADS | yes |

| | | | | |
|---|---|---|---|---|
| Ph43 | MKIAYARVSS | PRO9006_RS33035 | MSKPNGSARSREFTVPI | no |
| Sm18 | MITTNKVAIY | B0686_RS10845 | MKIAKKKWEPQIVNIMA | yes |
| Cd16 | MKQQIYNTA | DCP99_RS09010 | MNCLERFYRKGVCVIAD | yes |
| Pf80 | MKQAISYVRI | DOZ69_RS13980 | MQLIYSLNLKPQNNRDR | yes |
| Bs46 | MEDSSNKSV | BSL056_RS20850 | MNINKIRSFLYKTSKYLG | no |
| Rb27 | MQEHSPSNH | DUE16_RS19475 | MPKQQHANHRGNNDN | yes |
| Sa51 | MKGKIALYSR | DX957_RS00320 | MLKHQIINLIQEKREGSY | no |
| Bc30 | MTVGIYIRVS | IC1_RS37815 | MERTVLDNPPSEETKLR | yes |
| Cd04 | MNNRIDAIYA | A4Y97_RS04300 | MERKGDFISNNIRYKKE | yes |
| Sa34 | MNKVAIYVR | BJT53_RS05035 | MTHTKFNIITILPAGSKLK | yes |
| Pf15 | MRSAIPYIRF | AYO_RS35500 | MNYIVVHKPSQLIQKVIT | no |
| Ps45 | MKQAISYIRF | CCL18_RS05095 | MADVYNYRYGAIAQEFI | no |
| Ps45 | MKQAISYIRF | CCL18_RS04980 | MSYRKTSFEKHVDALHS | no |
| Sp56 | MSTSIPEESG | C550_RS27590 | MISALLDDVFDAELDAEI | yes |
| Dn29 | MERVLMHLR | DESNIDRAFT_RS0208480 | MKYDKTLKLGKTIIHIVNI | yes |
| Pc64 | MRVALYYRV | C747_RS0105930 | MTLPIDLDLLQKRKAIPVI | yes |
| Pc64 | MRVALYYRV | C747_RS0106115 | MSLNKLIIDTLKPIGVPVA | yes |
| Cp36 | MKQLNIQSSS | CPC_RS17335 | MKIKERNPPIKEYVKQV | yes |
| Pc01 | MRAAIVRRV | CHH67_RS12140 | MEKSERFEQEKRGISSSI | yes |
| Enc9 | MTALYARLS | ADH76_RS34160 | MEWEPEIKDSEFVPQKR | yes |
| Int2 | MPIAPEFLSL | SCAB_RS30855 | MTREERHRILSPAEIADA | no |
| Int3# | MRKVAIYSR | SPY_RS09195 | MKNKKEQWTPKVHCFR | no |
| Int6* | MAKKPKAKV | BMULJ_RS23410 | MTKKQSKHAEPTTDMQ | no |
| Int6* | MAKKPKAKV | BMULJ_RS23265 | MSKRPWQKWSEQEEA | no |
| Int10 | MITTNKVAIY | SP670_RS10955 | MNKRQRKKKILNGLNKE | no |
| Int10 | MITTNKVAIY | SP670_RS13965 | MKTVKKEWEPRIVNIM | no |
| Int13 | MAVGIYIRVS | BCER98_RS22635 | MKEIVVKPPQIPERTKRE | no |
| Int14 | MTVGIYIRVS | LIN_RS16360 | MSKVCDKENAVKVEIVY | no |
| Int15 | MKAAIYIRVS | LMOSLCC2372_RS12335 | MNKVLVSANYEGYESK | no |
| Int17 | MRTNEHNFH | SERP_RS07900 | MERSSVQFSTDGHGVR | no |
| Int17 | MRTNEHNFH | SERP_RS07570 | MSDKSKRTFNRWTSSE | no |
| Int21* | MRNKVAIYVI | SMI_RS10915 | MKVTVYAYGRKLEPDEE | no |
| Int22 | MKVATYVRV | GEOTH_RS21425 | MKKQKRVIKPQEVVVR | no |
| Int26* | MIAAIYSRKSI | CLI_RS20140 | MGFKVVVNYPTTEEGKI | no |
| Int29 | MKTAIYLRKS | BCERKBAB4_RS18200 | MPSTNMAVPTDQSHKH | no |
| Int30 | MYRPESLDV | GH772_RS27795 | MDSKTFKSGRVTIVIHSN | no |
| Int30 | MYRPESLDV | GH772_RS27775 | MSVIKQTSSDKDTKTIKS | no |
| Int32 | MDPQHKPTR | RHA1_RS50680 | MKDTFDRLPASWRREIF | no |
| Int33 | MKAIAIYARK | CLM_RS21070 | MAKIKKIIVNYPEDPKVM | no |
| Int33 | MKAIAIYARK | CLM_RS21065 | MAKGAEIKKVTIKIPEDT | no |
